# Trial-by-trial detection of cognitive events in neural time-series

**DOI:** 10.1101/2024.02.13.580102

**Authors:** Gabriel Weindel, Leendert van Maanen, Jelmer P. Borst

## Abstract

Measuring the time-course of neural events that make up cognitive processing is crucial to understand the relation between brain and behavior. To this aim, we formulated a method to discover a trial-wise sequence of events in multivariate neural signals such as electro- or magneto-encephalograpic (E/MEG) recordings. This sequence of events is assumed to be represented by multivariate patterns in neural time-series, with the by-trial inter-event intervals following probability distributions. By estimating event-specific multivariate patterns, and between-event time interval distributions, the method allows to recover the by-trial location of brain responses. We demonstrate the properties and robustness of this hidden multivariate pattern (HMP) method through simulations, including robustness to low signal-to-noise ratio, as typically observed in EEG recordings. The applicability of HMP is illustrated using three previously published datasets. We show how HMP provides, for any experiment or condition, an estimate of the number of events, the sensors contributing to each event (e.g. EEG scalp topography), and the intervals between each event. Traditional exploration of tasks’ cognitive structures and electrophysiological analyses can thus be enhanced by HMP estimates.

## 1 Introduction

The speed of information processing in the brain has connected psychology and physiology since Von Helmholtz assumed the existence of processing steps in the reaction time (RT) to measure neural transmission velocity in humans (von Helmholtz, 1850). Since then, researchers have investigated the nature and speed of the different information processing operations – or cognitive processes. Classical mental chronometry (Posner, 1978) tackled this problem by relying on experimental manipulations. As one of the earliest examples, the subtractive method developed by Franciscus C. Donders (1868) takes as an indirect measure of a cognitive process’s duration the difference in mean RT from two tasks that are assumed to differ in the insertion of that particular process. For example, the duration of a response selection process could be inferred from the RT difference between a task with response selection, and the same task without response selection.

The RT as a composite measure of the time needed for different cognitive processes to complete remains a shared assumption between the main theories of mental chronometry (Luce, 1986). This assumption is further fueled by the repeated observation that RT distributions do not obey any of the known and commonly found statistical distributions (Christie & Luce, 1956; Noorani & Carpenter, 2016). Therefore, considerable effort has been devoted to decomposing RT distributions into different underlying distributions, for example by a Fourier transform approach (Green, 1971; Smith, 1990), mixture modelling (Christie & Luce, 1956; Van Maanen et al., 2014), cognitive (e.g., Anderson et al., 2018; Tenison et al., 2016; Van Maanen et al., 2009), mathematical models (e.g. Anders et al., 2016; Brown & Heathcote, 2005; Ratcliff, 1978; Smith, 1995), and combinations (e.g., van der Velde et al., 2022). While all these applications have extended our understanding of several aspects of human cognition through mental chronometry, the reliance on RT alone limits the ability to decide between competing hypotheses (Forstmann et al., 2011).

### 1.1 Mental chronometry and EEG

To extract the nature and unfolding of cognitive processes, neural recordings with high temporal resolution such as electro-encephalography (EEG), magneto-encephalography (MEG) or intra-cranial recordings would stand as strong candidates. This has led Meyer et al. (1988) to call for a modern mental chronometry approach where physiological recordings inform our understanding of cognitive processes involved in the RT and vice-versa (see for example Kelly & O’Connell, 2013; J. Miller et al., 1999; Purcell et al., 2010). However, measuring the location of cognitive events in physiological signals is complicated by their low signal-to-noise ratio, as relevant activities are usually several orders of magnitude weaker than the background noise of these measures.

Most research efforts have therefore relied on signals averaged over many trials such as peri-stimulus time histograms, event related potentials (ERP), etc. Unfortunately, averaging continuous time series across trials is well known to hide and distort single-trial effects (e.g. Burle et al., 2008; Callaway et al., 1984; Luck, 2005). These distortions arise because of the time variation in the by-trial generators of these average curves, making inferences about the timing of the generators complex (Luck, 2005). These distortions are exacerbated when comparing condition-averaged signals: if the process or component time jitter distribution differs between conditions – as could be expected from behavioral (Noorani & Carpenter, 2016; Ratcliff, 1978; Weindel et al., 2021) and electrophysiological studies (O’connell et al., 2012; Smulders et al., 1994) – then both the peak amplitude and latency of the averaged signal will be different across conditions (Mouraux & Iannetti, 2008).

Additionally, when researchers are interested in several components to describe a task, the temporal overlap of components close to each other also distorts the average of the components (Luck, 2005). Lastly, if one assumes that components in an experimental task are sequential to one another (*e*.*g*., if the time of the response-related potential is causally affected by the time of the visual-related potentials), then the time jitter in the later component will contain jitter from the previous components. This has prompted researchers to use multiple events to which to time-lock the signal, for example by using either stimulus onset, response onset or other events such as electro-myographical onset (see Burle et al., 2004, for a comparison of the three methods). However, most psychological processes involved in a task will not have a relevant external index to time-lock physiological analysis on.

For true modern mental chronometry it is therefore necessary to have single-trial descriptions of neural time series. Several single-trial time measurements of physiological events have been suggested. Nunez et al. (2019), for example, used EEG to measure a single-trial visual encoding time and related it to experimental manipulations. Kutas et al. (1977) used a pattern search for an EEG component – the P3, associated with stimulus evaluation – to obtain by-trial measurements of the timing of the component based on properties of the average ERP (Woody, 1967). Furthermore, electro-myography was used to capture motor execution time and deepen our understanding of the link between the decision and the motor system (Botwinick & Thompson, 1966; Burle et al., 2002; Weindel et al., 2021). All these applications show that the addition of external recordings is a valuable addition for in-depth exploration of mental processing. Nevertheless, these applications only isolated one processing component from the others, while almost any cognitive task consists of multiple processing steps.

### 1.2 Single-trial estimation of cognitive events

The two main theories of event-related potential generation: the phasic burst (Makeig, 2002) and the synchronized oscillations theories (Basar, 1980, as cited by Anderson et al., 2016) predict that a significant cognitive event is expected to be visible as a transient change in an EEG timeseries. Anderson et al. (2016) proposed a method using a time-based regime-switching model that assumes that these transient changes are multivariate patterns repeating across all trials in a sequential order. In addition to the sequential appearance of the events, the method assumes that the repetition across trials of these peaks follows a specific probability distributions. By using this method on EEG or MEG data, and therefore estimating these patterns and their distributions, one obtains a time location in each epoch for each of these putative sequential events. Hence, contrary to ERP averaging, or pattern search based on properties from the averaged ERP (Kutas et al., 1977; Smulders et al., 1994), the method by Anderson et al. (2016) uses expected time, sequentiality of the patterns and channel contribution to uncover single-trial EEG or MEG events across the whole time-series defined by the RT.

This idea of sequential by-trial time varying events has been applied to a wide range of tasks, ranging from simple perceptual (Van Maanen et al., 2021) or lexical decision tasks (Berberyan, van Rijn, & Borst, 2021) to more complex tasks, such as tasks involving visual working memory (Zhang et al., 2018), associative recognition (Anderson et al., 2016; Portoles et al., 2022), and mathematical problem solving (Anderson et al., 2018; Groeneweg et al., 2021). Each of those studies provided new insights on the nature and latencies of cognitive events of the analyzed tasks. For example, in the original paper, Anderson et al. (2016) linked the decomposed RT to a cognitive model of the task. Van Maanen et al. (2021) showed how to separate a decision stage from the other stages and could therefore relate the single-trial decision stage times to an evidence accumulation model (see also Berberyan, van Maanen, et al., 2021). Berberyan, van Rijn, and Borst (2021) showed that word frequency and lexicality effects are driven by a single cognitive processing stage.

In the present study we aim to formulate a generalized and multi-purpose model of these sequential by-trial multi-variate patterns based on the work by Anderson et al. (2016). We define a method with flexible definition of the single-trial pattern and their expected interval distributions. Furthermore we create a fitting routine to infer the number of events that are needed to account for the M/EEG data instead of assuming a given number a priori. In the remainder of this paper, we will showcase the use of the HMP method both in simulated and real EEG data, after providing a formal description of the method. All analyses that we report throughout the paper are performed with the accompanying Python package (see the online repository, https://github.com/gweindel/hmp, including a set of tutorials), illustrating the kind of results that are readily accessible to the interested researcher.

## 2 The hidden multivariate pattern method

The hidden multivariate pattern method or HMP describes a multivariate time-series (e.g., EEG or MEG channels as in Figure 1a), observed from time 0 (e.g., stimulus presentation) to an end time *T* (e.g., the moment of the response, defining the RT), as a linear succession of a given number of events (Figure 1b). These trial-recurrent sequential events are each assumed to be represented by a pattern with a given duration (Figure 1c) produced by a specific mixture of channels (Figure 1b and d, see Section 2.1.1).

**Figure 1.**
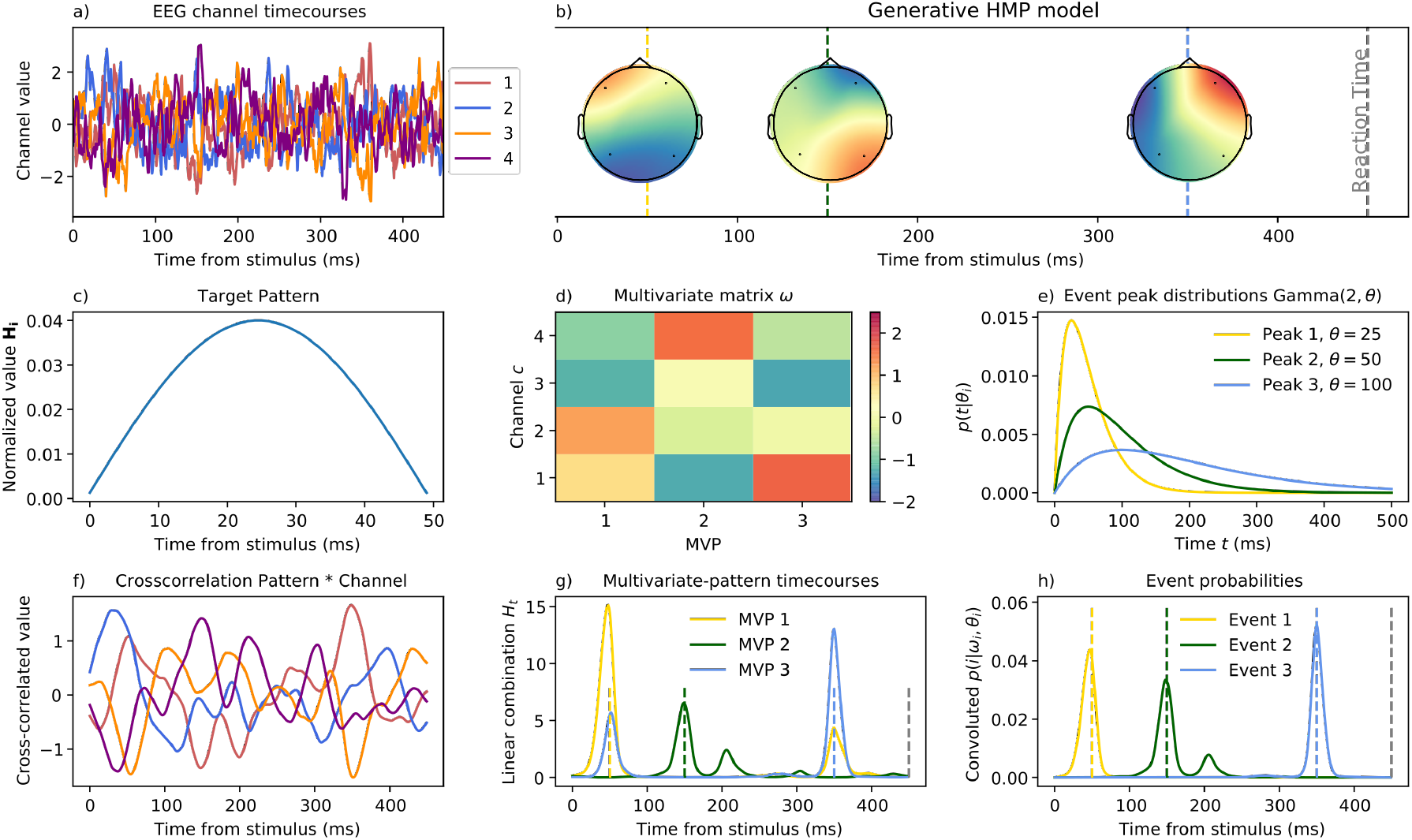
Example of the estimation for a three-event HMP model on simulated data for four EEG channels. Panel a) shows the simulated time-series, locked to stimulus onset time, for the four EEG channels on a trial with an RT of 450 ms. These time-series contain the three events depicted in b) whose peak are at times 50, 150 and 350. The pattern is a 50-ms monophasic pulse (half-sine) represented in panel c). The multivariate representation of, i.e. the contribution of each electrode to, each of the three events are color-coded in panel b) and d). Panel e) shows the probability distributions of the peak of each event relative to the previous event (stimulus in the case of the first event) therefore indicating how likely each event is after a given time following the previous event. The contributions and the distributions for each event are typically estimated from the data. In order to estimate an HMP model, the channel data are first cross-correlated to the expected pattern resulting in the time-series displayed in f). These cross-correlated signals associated with the mixing matrix displayed in d) give the three multivariate patterns (MVP) time-series for each of the three expected events (g) where higher amplitude indicates higher likelihood of the presence of the multivariate-pattern. By imposing the distributions in e) and the assumption of sequentiality, we can estimate the probability of the peak of each event knowing its multivariate representation and expected peak time, and conditioned to the probability of peaks of the preceding events. The sequentiality and time-distribution constrain can be seen by comparing panel g and h, in panel g the multivariate pattern time-series of the first and last event (yellow and blue) share the same times, but once the convolution is applied, the evidence for the first event is only found around 50 ms and the evidence for the last event is only found around 350 ms. In HMP the parameters displayed in d) and e) are estimated from the data and the resulting probabilities in h) are used as proxy for single-trial event position.

The peaks of the multivariate patterns are assumed to be sequential to each other and the distances in time between them follow probability distributions of which the parameters are estimated from the data for each event (Figure 1e, see Section 2.1.2). The sequentiality of the events implies that the expected peak time of the *n*th event depends on the expected peak time of all previous events. This sequentiality implies a fairly complex problem that is resolved using dynamical programming (see Section 2.2.1).

In theory, the method can be applied to any defined time interval and any type of measurement, however, the current presentation (and Python implementation) focuses on the time interval defined by the RT duration from a subject using EEG or MEG data. The application of the method results in an estimation of the probabilities for each of the detected events at each sample for each trial in the data used for the estimation (Figure 1h, Section 2.2.4). These probabilities can be used for follow-up analyses, which we will illustrate in Sections 3 and 4.

### 2.1 HMP Concepts

#### 2.1.1 Pattern analysis

An HMP model assumes that events are represented by patterns in the data, such as the half-sine, or ‘bump’ (Figure 1c), proposed by Anderson et al. (2016) and based on the main theories of ERP generation by Makeig, 2002 and Basar (1980, cited by Anderson et al., 2016). Once a template **H** is expressed with a certain duration, it is cross-correlated to the signal of each channel *c* to reflect the similarity of that channel at each time point *t* with the template (an example definition of **H** can be found in section 2.4). The dot product between each sample *s* at time *t* with the template of length *L* is computed :

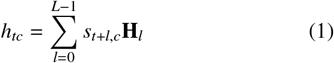

After this transformation, the sample *h*_*tc*_ reflects the cross-correlation of the template with the signal at time *t* for a given channel *c* (Figure 1f). This results in a measure of similarity between the template **H** and the sensor data *s*_*tc*_.

In the case of several sensors that record brain activity, the multivariate patterns are defined as a linear combination of the *C* channels. The magnitude of the contribution, *ω*, is defined as a matrix of size *C* × *I* with one value for each sensor *c* and event *i* (Figure 1d). The match between the channels and the event *i* at time *t* is written as *H*_*ti*_ (Figure 1g):

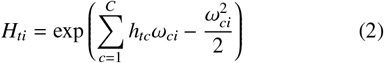

The exponential ensures that we keep the match positive and the subtraction controls the relative contribution of the pattern presence *H*_*ti*_ with the expected times in the next section (see Appendix 1 from Anderson et al., 2016). When *I >* 1, *ω* is a vector of size *c* for each event *i*.

#### 2.1.2 Inter-event time interval

In HMP, the peak time of event *i* depends on the peak time of the event *i −* 1, expressing their expected time dependency. This dependency has to be expressed by a distribution on positive real numbers for which the mean peak time can be derived from one parameter (*e.g*., a gamma distribution with a shape fixed to 2 as in Figure 1e). For the following, we write *p*(*t*|*θ*) as the probability density function of the peak time *t* given by a parameter *θ, g*(*t*; *θ*), divided by the sum of its elements:

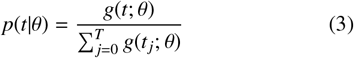

In the case of only one event (*I* = 1), the overall duration *T* of a trial (i.e., the RT) is partitioned into two intervals with durations *t* and *T*−*t*. Therefore, only one distribution of event peak needs to be estimated. The probability of the event peak *i* happening at time *t, p*(*i* = *t*), is given by the likelihood of the multivariate pattern associated with *i* expressed by cross-correlation *H*_*t*_ across all sensors, modified by the probability that this event occurs at time *t*−1 and the time of the response occurs after time *T*−*t* given *θ* (e.g. the yellow curve in Figure 1h):

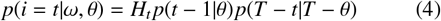

In the case of *I >* 1 one *θ* is estimated for each event.

### 2.2 Estimation of an *I*-event HMP

The number of free parameters for an *I*-event HMP is *I* × *C* + *I* + 1 with *I* + 1 peak time parameters *θ* and *I* × *C* magnitudes *ω* of the channel contributions. When *I >* 1, to compute the probabilities of peak time for event *i* when *i >* 1, one needs to integrate all possible time locations for the previous *i* − 1 events. Given the complexity of such computation the method relies on dynamic programming using the Baum-Welsh algorithm (Visser, 2011).

#### 2.2.1 Inference

The estimation relies on an adapted version of the forward-backward algorithm usually used in hidden semi-Markov models (see Yu, 2010, for a formal description of the algorithm). The forward variable *α* at a given time *t* tracks the probability of an event given past sequences from 1 : *t*. The backward variable *β* accounts for the probabilities of coming observations given the presence of an event. Multiplied together, *α* and *β* give the likelihood of a an event *i* being located at time point *t*.

For an *I*-event model, assuming that the parameters *θ*_*i*_ and *ω*_*i*_ are known, the forward variable *α* at time *t* for a given event *i* is:

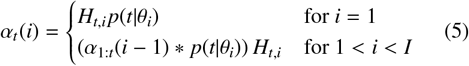

In Equation 5, (*α*_1:*t*_(*i* 1) *p*(*t θ*_*i*_)) is the convolution between the value of the forward variable for the previous event and the expected time before the next event. This convolution allows us to account for the different time points at which the previous event could have happened and captures the core assumption of event sequentiality in an HMP model. This value is then multiplied with the exponentiated sum of the cross-correlation magnitudes (*H*_*i*_) for the HMP signaling the next event.

The value of the backward variable *β* is:

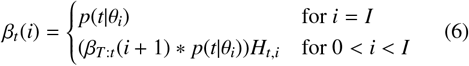

The likelihood of a particular event *i* time location for time point *t* is then:

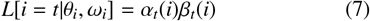

Because of the cross-correlation of the signal to the pattern, one needs to ensure that the samples within an event cannot be associated with a new event. For this reason, in the current estimation procedure of an HMP model, we chose to censor in Equation 3 the samples within the width of the pattern during the EM phase. This imposes the constraint that the to-be-detected event peak time following another has a minimum average interval of the expected pattern duration. At the end of the EM phase this constraint is relaxed allowing thus to estimate single-trial times even for event intervals shorter than this censoring. This censoring is not strictly necessary but greatly simplifies the estimation of HMP with distinct events (see Appendix B and the Discussion section).

From Equation 7, we can derive the probability of an event *i* at time *t* after event *i*−1 by normalizing over all likelihoods across samples:

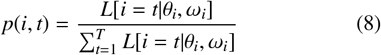

Figure 1 represents an example of the estimation steps (bottom row) on simulated EEG data for a three HMP model (top row), given the parametrization described in Section 2.4 (middle row).

#### 2.2.2 Expectation maximization

In most cases, parameters *θ*_*i*_ and *ω*_*i*_ are unknown and need to be estimated from the data. For brevity, we write *λ* as being the set of the *I* + 1 scale parameters *θ* and *I* × *C* magnitudes *ω*. We estimate the joint vector of parameters *λ* = (*θ, ω*) that best describes the data through expectation maximization (EM, Rabiner, 1989). We first make a proposal on the values of *λ* and compute the log-likelihood of the events summed over the sequence 1 : *T*

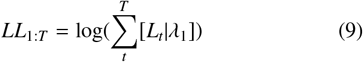

The EM algorithm takes a first proposal to compute the event probabilities of Equation 8 and uses these to compute better parameters *λ*^*^. In the case of the intervals between two events *i* and *i* − 1 (time from sequence start for *i* = 1, e.g. stimulus), *θ*^*^ is computed from the probability-weighted average interval as given by the most likely position of each event, from which is subtracted the previous event position:

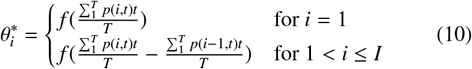

The function *f* to go from intervals between events to scale parameter 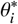 depends on the chosen distribution (see example for a gamma with shape 2 in Section 2.4).

The magnitudes of the channel contribution to the events (matrix in Figure 1d) are computed based on the average channels values at the event location times, scaled by the probability of the event:

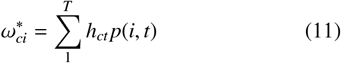

Equation 9 is then computed on the basis of the new set of parameters *λ*^*^. In our application, the EM algorithm stops as soon as the relative difference in log-likelihood between *λ*^*^ and *λ* increases less than a given tolerance *ϵ*:

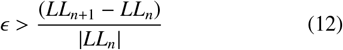

#### 2.2.3 Repeated measurement

With the exception to the case where no noise is present in the data (e.g. Figure 1), the identification of HMP events requires multiple trials. In this case, we assume that each trial is an independent new realization of the same event sequence. *T* is then the maximum duration of all fitted trials. The missing values in the sensor signals for trials shorter than *T* are imputed with a gain of 0 in Equation 1. EM Equations 10 and 11 are adapted to take the mean across all trials, and Equation 9 is then summed across all trials.

#### 2.2.4 From event probabilities to by-trial event times

When fitting an HMP to a time-series, researchers will usually be mainly interested in the probabilities computed from Equation 8 and displayed in Figure 1h. These posterior distributions capture for each event the certainty about the time location of an event based on the estimated model. From there several strategies are possible to attribute a single time to each event at each trial. One can either use the weighted average as in Equation 10 or any other method (mean, max) to turn the event probabilities into single time points to use in a follow-up analysis. In the present manuscript we take the maximal probability of each event as the optimal estimate of the true time points at which the peak of the event is located. Once the most likely time point of each event at each trial has been selected, we compute the values of the channel at the peak of the expected event. Using this method allows to average trials and obtain time based topographies within the RT as plotted in the top row of Figure 1 and as illustrated in the following sections. These times can also be used as starting point for additional analyses as done throughout the remainder of the manuscript.

### 2.3 Finding the number of events

How many events are present in a task is often part of the scientific question, we derived two ways of estimating how many events are needed to describe the recorded time-series.

#### 2.3.1 Cumulative fit

The first strategy to estimate the optimal number of events needed to account for the data, is to exploit the sensitivity of the EM algorithm to local maxima (Wu, 1983). That is, given an initial proposal of the time location parameter *θ*^*^, the EM algorithm finds the nearest local maximum. In our application, this entails that a one-event HMP model with a location parameter initialized at the first sample 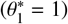 will converge (within a certain tolerance *ϵ*) to the log-likelihood local maximum of the first “true” event. In this procedure, the magnitudes are typically initialized at 0, although this is not strictly necessary. The first local maximum, being the first event, is thus parametrized by *λ*_1_ = (*θ*_1_, *ω*_1_).

After identifying the first event this way, we fit a two-event model in which the first event is initialized with the optimal parameters of the one-event HMP model (*θ*_1_, *ω*_1_), and the second event again with magnitudes of zero. The time sample following the most likely event time, as defined by Equation 8, is then used as proposal location parameter for event 2. This way, the EM algorithm will find the second local maximum. Next, a three-event HMP model is fit, initialized in the same way, and so on. This procedure continues until the average RT is used as proposal time location.

Once we cumulatively fit HMP to the data, a model with the *n* events obtained this way is fitted to the data, using the parameters associated with those events as initial proposals. This model fine tunes the location and magnitude of the optimal number of events that can be found in the data. Simulations and application to real data in the present manuscript show that this sequential estimation procedure performs well in recovering the exact number, location, and magnitudes of underlying events. If necessary, this can be followed by an LOOCV procedure, for example to test which events generalize over subjects (Anderson et al., 2016; Van Maanen et al., 2021).

#### 2.3.2 Alternative fitting strategies

The cumulative event fit method described above relies on the detection of local maxima where the EM algorithm converges. While this method allows to estimate an HMP model without specifying a number of events a priori, it does however make the assumption that the local maxima can systematically be found by cumulatively fitting HMP models. Alternatively, researchers can fit an HMP model with a prespecified number of events to explore different event number hypotheses. In this case however, one needs to ensure that the parameter space has been properly explored, for example by running several starting points for the parameters of HMP. Once an HMP with a given number of events has been estimated with enough certainty with respect to starting points, researchers can test for the generalization of each event by iteratively removing an event and testing the generalizability of the resulting *n* − 1 HMP with the LOOCV method (see Borst & Anderson, 2024, for a detailed explanation).

### 2.4 Example HMP parameterization

The general formulation described above ought to be parameterized according to a hypothesis about the pattern of the underlying events and on the overall shape of the probability distribution used between events. In the simulated and application to real data sections we use a half-sine pulse with a duration of 50ms as template and a gamma distribution with a shape of 2 as described by Anderson et al. (2016). To match the half sine-wave frequency *f* (10 Hz in this example) and the sampling frequency of the signal (*f*_*s*_), we construct a vector **V** of length 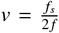 with values 0, 1, …;, *v* − 1. Values within **V** are adjusted to the signal sampling frequency (adjusted to ms) by multiplying by 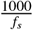 and adding 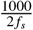 to estimate the sine-wave value half-way between samples. The sine wave associated with *f* is then sampled accordingly:

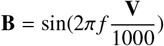

The sum of its squared elements then normalizes the resulting vector:

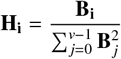

**H** is then the normalized template of a half-sine wave with duration given by *f* (Figure 1c). For the probability distribution we choose a gamma with a shape fixed to 2 and a scale estimated from the data. To resolve Equation 10 in the estimation procedure, *θ* is calculated from the average event time peak of a starting point or a previous iteration of the EM algorithm as 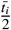, where 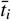 is the average interval for event *i* relative to the peak of the previous event. Conversely, the average time interval 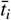 in the case of a gamma with a shape of 2 is given by 2*θ*_*i*_.

## 3 Simulations

To illustrate properties of the method we simulated EEG data and tested the recovery of the ground truth by HMP. All the simulations rely on the MNE Python package for analyzing M/EEG (version 1.6.1; Gramfort et al., 2014) and use MNE’s sample participant channel positions, between electrode variance-covariance matrix and forward model. In these simulations, events are defined as the activation of a source in the brain, as defined by the source space, with projection to the electrode space as defined by the forward model. The signal is simulated continuously with epoch start triggers inserted at different times within the length of the simulated signal. For each of these thereby defined sequence/trial starts, except when mentioned otherwise, events are inserted with their position relative to the previous event (sequence start for the first simulated event). The end of a trial is defined as a last event (without any source activation) inserted randomly following the before last event. Figure 2a1 provides an example of four HMP events and a “behavioral response” event on three simulated trials, without any noise. The vertical lines indicate the event location, the time series represent the amplitude of the signal for a selected electrodes (EEG 038).

**Figure 2.**
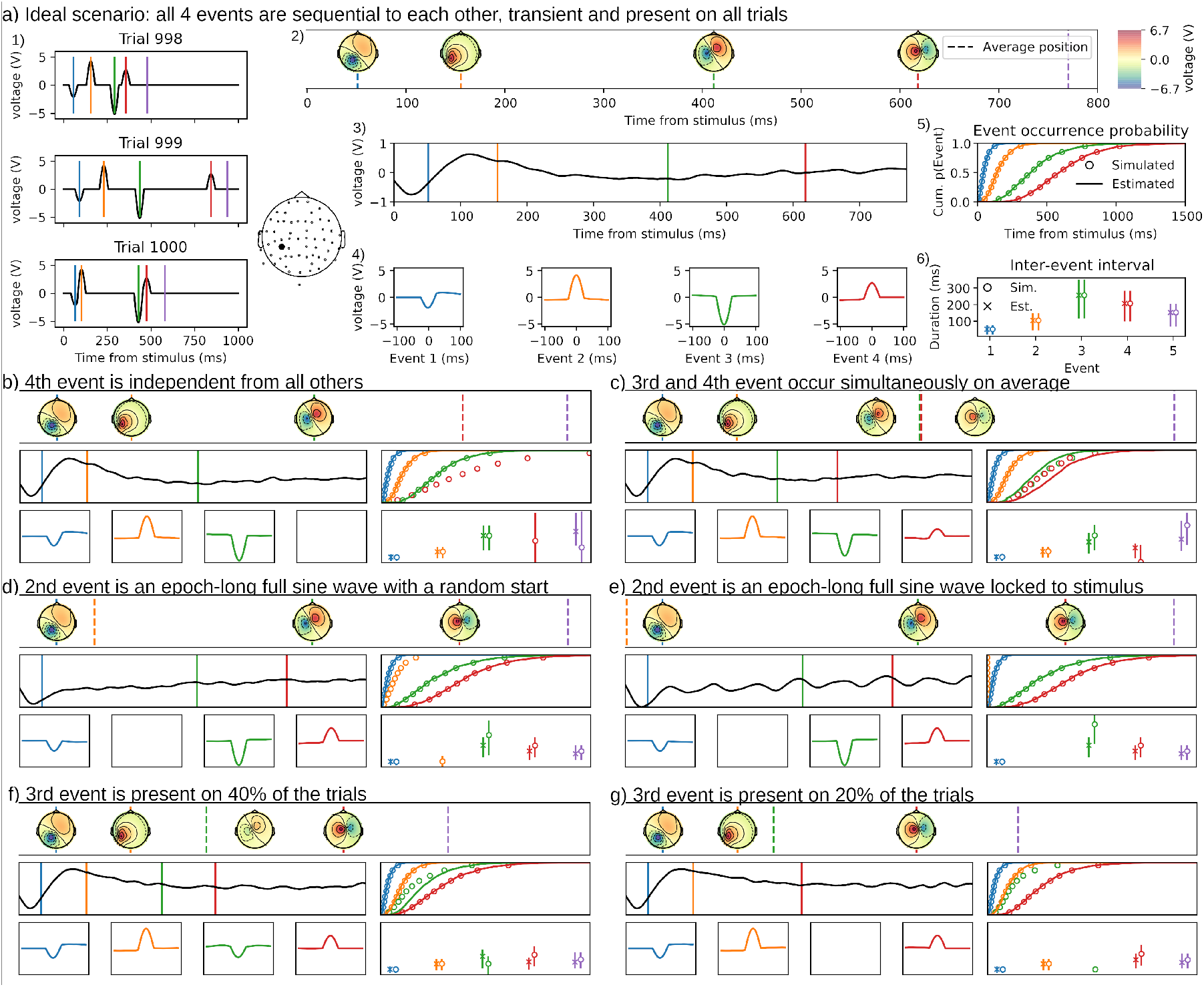
a) Ideal simulation illustrating a perfect recovery with HMP a.1) shows the simulated signal for 3 example trials for one electrode whose spatial location is depicted next to a.1). The vertical lines indicate at which times, per-trial, the 5 events (4 peak source activity and a response time in purple) occurred. a.2) Represents the, by-trial time aligned, averaged topographies at the trial-averaged times of each event. Dashed lines indicate the simulated trial-averaged event times. a.3) Average ERP for the electrode centered on the stimulus, the colored lines are the average time of the estimated events. a.4) average electrode activity centered around the by-trial peak of each event. a.5) Cumulative probability of event occurrence, excluding the response, over time with the deciles (5th to 95th percentile) for the simulated distributions (circles) and the estimated empirical distributions (plain lines). This figure represents how far the estimated event positions are from the simulated ones. a.6) distributions (mean, 1st and 3rd quantile) of difference in time between the by-trial most-likely location of the successive events for simulated (circles) and estimated (cross) intervals. This figure captures how the by-trial times between the events are recovered. Simulations from b) to f) all use the same seed and simulation structure as a) except on one point detailed for each panel hereafter. The figures are all on the same scale as panel a). b) The location of the last event before the response is independent of the other events. This means that it is not strictly following the 3rd event nor strictly preceding the response. In this case the event is not detected by HMP. c) Events 3 and 4 are occurring simultaneously. HMP detects both but wrongly estimates the times of each. d and e) 2nd event is turned into an oscillatory activation (pure 10Hz sine-wave) lasting all the epoch either with a random start (d) or a time-locked to the simulated stimulus onset (e). In both cases the oscillatory event is not detected but the others are detected. f and g) the 3rd event is only present on some trials, either 40% of the trials (f) or 20% of the trials (g). If the proportion of trials displaying the events is too low, HMP will not detect the event.

In this simulation section, we first simulated EEG data in varying scenarios but without any noise added to the signal. This section serves as both an illustration of the method and its assumptions, and the consequences of a failure to meet these assumptions even in the case of a perfect signal. Second, we tested the robustness of HMP to noise under several signal-to-noise ratios to illustrate its potential usefulness for signal-to-noise ratios closer to what can be observed in actual EEG and other neural time-series data.

### 3.1 Method

#### 3.1.1 Defining simulation parameters

##### 3.1.1.1 HMP event simulation

In the simulations, we choose to implement HMP events as the activation of a source in the MNE sample participant’s source space that yields a multivariate pattern (Figure 1c and d combined) at the channel level (59 EEG electrodes) following the forward model provided with the sample participant. An HMP event is then simulated by the activation of a source following a pattern (half-sine) with a given time duration added to the signal. These thereby defined events are then inserted at different times in each simulated trial sampled from the chosen probability distribution for the inter-events intervals. For each trial, the last simulated event defines the behavioral response event and therefore the simulated RT. The number of HMP events in each trial is the number of sources that were simulated in the time-series, excluding the last source used for the simulation of a behavioral response (last lines in all simulated HMP models representations).

##### 3.1.1.2 HMP estimation

The simulated datasets were epoched from simulated trial start to simulated response event on each trial. Each trial was then centered and we performed a dimension reduction on the data by applying a PCA, common to all trials, on the average variance-covariance matrix of the electrodes (Cohen, 2014) and keeping the 5 first components (therefore applying HMP on 5 virtual channels). The resulting virtual channels were z-scored for each epoch. An HMP model was then initialized as described in Section 2.4 with a target pattern of 50ms, a gamma with a shape of 2 and a scale free to vary (see Appendices B, and C for alternative parametrizations). The tolerance for the expectation-maximization (Equation 12) was set to 10^−4^ (a stricter criterion provides roughly the same results but estimation takes longer). We then use the cumulative event fitting method described in Section 2.3.1 to estimate the number, by-trial times and topographies for each HMP event.

### 3.2 Illustration of HMP properties

For the purpose of illustrating HMP we generated 7 simulations with 1000 trials without any noise in the signal and with a sampling frequency of 1000 Hz. The first simulation is the idealized simulation in the sense that this simulation is in accordance with all the core assumptions of HMP: all events are sequential to each other, implemented as transient events present in all trials. Each of these core assumptions were then violated one-by-one, with two different violation types, in the subsequent simulations. The resulting 6 degraded datasets are used to illustrate the consequences of a failure to meet these core assumptions.

#### 3.2.1 Idealized simulation

This simulation is constructed by inserting four sequential 50-ms wide HMP events (see the average topographies and times in Figure 2a2) between each trial’s start and simulated response event. The times of each event are sampled from gamma distributions, all with a shape of 2, with scales of 25, 50, 125, 100 respectively from each previous event. The between-event intervals are therefore 50, 100, 250 and 200 ms on average (see Section 2.4). The time from the last event to the response is sampled from a gamma distribution with shape 2 and scale of 75 (thus 150ms). Three simulated trials using this procedure are presented in Figure 2a1 for one electrode (EEG 038).

As can be seen in Figure 2a2, HMP recovers the four events at the average simulated times. To show how the by-trial positions can be informative in this simulated case, we plot the average ERP for a selected electrode (EEG 038) that display a maximal response for all the four events. The average stimulus-locked ERP (Figure 2a3) displays the classical properties of real-data average ERP, with an overall curve much more spread out and of lower amplitude than the single-trial simulated event displayed in Figure 2a1 for the same electrode. Given that no noise was simulated, the discrepancy between the stimulus-locked ERP and the underlying single-trial signal, results solely from the between trial time jitter of the events as well as overlap between components.

By fitting an HMP it is possible to recover the by-trial event time-jitter exemplified in Figure 2a1 and time-lock the ERP to the detected single-trial peak times of each event. This procedure is illustrated in Figure 2a4 where the average electrode activity around each event is calculated after the data have been time-locked to the by-trial time of each event. With this representation we clearly see all events independent of their position with respect to stimulus onset, with an average waveform congruent in shape and voltage peak with the simulated 50 ms event. Figure 2a5 and a6 shows the time properties of the estimated HMP events. Figure 2a5 represents the event occurrence probability in time (i.e. similar to Figure 1h but averaged over trials and transformed to a cumulative sum) and shows the match between the estimated probabilities of the HMP events (solid lines) vs the time at which each by-trial event was simulated with respect to stimulus onset (circles representing percentiles from 5 to 95% in steps of 10). From this panel we can evaluate how HMP recovered the position of the event in each epoch. Figure 2a6 illustrates the 1st and 3rd quantiles (vertical lines) and the mean (circle) of the inter-event intervals distribution, using the most likely position of each event (as defined in Section 2.2.4). From these intervals we can evaluate that the recovery of the average simulated intervals of 50, 100, 250, 200 and 150ms between each event is accurate.

Having established that, in an idealized simulation, HMP correctly recovers the events, we now turn to simulations in which core assumptions of HMP are violated.

#### 3.2.2 Violating the sequentiality assumption

##### 3.2.2.1 Non-sequential event

To violate the sequentiality assumption we first simulated the 4th event as being linked to stimulus onset (with the same expected mean position in the epoch) instead of following event 3 and being followed by event 5 (i.e. the response). In this simulation, the response event is driven by the 3rd event with an expected interval of 200 + 150 ms to obtain an expected RT at the same position as in the idealized simulation. Because the sequentiality assumption is violated, we observe that, despite no noise, the 4th event is missed by HMP but the other, sequential, events are found with the same accuracy as in the idealized simulation. This is reflected by the missing red event in all panels of Figure 2b. The reason why the non-sequential event is not found is because HMP dynamically locates each event based on all the other ones. If, as in this case in this simulation, an event’s location is independent from the other, the neighboring event locations don’t inform the location of that independent event as it can occur before or after those events on any trial.

##### 3.2.2.2 Simultaneous events

If instead we simulate the 4th event dependent on the 2nd event and on average at the same time as the 3rd event (e.g. one brain potential leading to two independent but simultaneous events) then HMP recovers each event but cannot recover the timings accurately as they overlap in time. This is reflected in the event occurrence probabilities and the interval estimation where the estimation of the time of the 3rd event is shorter than simulated, and the time of event 4 is longer than simulated. Furthermore the event centered ERP for the events 3 and 4 also reflect this time misalignment as the HMP centered ERPs are of smaller amplitude than in the idealized simulation. Interestingly, despite simultaneous on average, the 3rd and 4th events are close to their ideal topography counterpart, showing that HMP separated these as two different events instead of generating averages of them. This example illustrates that two events can be simultaneous on some trials yet be recovered as different events as long as the interval between both varies over trials.

In summary for the sequentiality assumption, if a topography repeated across trials appears tied to the stimulus but not the the other events it will not be detected nor be a problem for the estimation of the other, sequential, events. If two events are the consequence of a same previous event and have the same timing, then both are recovered but a time misestimation can occur.

#### 3.2.3 Violating the assumption of transient events

In this simulation we simulated the 2nd event as being oscillatory instead of being a transient half-sine. This is achieved by changing the 2nd event from a 50ms activation to a 10Hz pure sine-wave (alpha wave) from event start to the end of each epoch. In this case, event 2 is missed by HMP, independent of whether the event has a random phase and start in the epoch (Figure 2d) or whether it is time-locked to the stimulus (Figure 2e). Reassuringly however, despite our target pattern being exactly half a period of the simulated oscillatory event, all other event estimations are unaffected by these simulated oscillations. In the case of the stimulus phase-locked alpha it is particularly interesting to observe the disconnection between the stimulus centered ERP (Figure 2e, middle panel), largely contaminated by the oscillations, and the HMP centered ERPs (Figure 2e, bottom panel) that do not display a contamination compared to the same plot in the idealized simulation (Figure 2a4).

In summary for the assumption of a transient event, oscillatory events are not detected but also, in the absence of noise, do not represent a competition for the estimation of the true transient events.

#### 3.2.4 Violating the trial presence assumption

Another realistic scenario could be that an event is only present on a proportion of trials. This could for example occur when some trials display a particular cognitive process, but others do not due to different strategies (Archambeau et al., 2023; Groeneweg et al., 2021). We simulate two cases where the 3rd event is simulated only in 40% (Figure 2f) or 20% (Figure 2g) of all trials. As can be seen in the corresponding panels, if the event is present in at least 40% of the trials, HMP recovers the rare event. Nevertheless the estimates of the event timing are slightly off. The disconnection arises from the fact that on 60% of the trials the HMP detected event is not present but HMP assumes that events are present on all trials, thus, the model will place the event on the most likely position given the previous event time and the estimated time distribution for this rare event. In the case where this event is simulated in 20% of the trials, the event is simply missing in the estimation, but the estimation of the remaining events is barely influenced (only in the 20% of the trials with that extra event will the timing of the next event be off compared to the simulated times).

In summary it appears that HMP is robust to violating the assumption that the events are present on all trials. If an event is present in enough trials (40% in the case of this simulation without noise) the event will be recovered, but in this case the estimated interval between events is less well recovered as the event time is estimated by HMP at each trials, including trial where it is absent.

Overall, all these illustrations serve to show the core assumptions of HMP of transient sequential events repeated across trials. HMPs core assumptions are useful to locate events in noisy recordings. Whereas these simulations were performed on a noiseless signal, the quality of the estimation in an HMP mode on real data (and therefore also the robustness to the core assumption violation) will depend on the signal-to-noise ratio. We will explore this in the following section. Ancillary assumptions related to the choice of patterns and of the time distributions are explored in Appendices B and C.

### 3.3 Signal to noise ratio simulation

In this simulation we investigated the effect of different signal to noise ratios (SNR) on HMP estimations. We simulated data as in the ideal case of Section 3.2 for 3 and 5 events. The by-trial interval between the peak of the events (and the interval from the last event to the behavioral response) where randomly sampled from a gamma with a shape of 2 and a per-interval scale itself drawn randomly from a uniform distribution between 25 and 150 (thus an average inter-event interval between 50 and 300 ms). The noise was drawn from a multivariate Gaussian with variances of 0.2 *µV* and covariances among electrodes given by MNE’s sample participant. The generated noise was filtered with an infinite impulse filter (IIR, with values 0.2, -0.2, 0.04 respectively for the feedforward, feedback and gain coefficients) to increase the resemblance to actual EEG noise. The SNR was determined using the average of the squared voltage value across trials at the peak of each simulated event divided by the variance across trials at the same time-points.

To test the impact of different noise levels we selected three source amplitudes corresponding roughly to an average SNR of 1/10, 1/2 and 1/1. Figure 3a shows the single-trial activities associated with all 3 SNR levels for three sources in a trial. We also illustrate the SNR levels with classical ERPs in Figure 3b for two selected electrodes, suggesting that typical EEG data has a higher SNR than all three levels simulated here.

**Figure 3.**
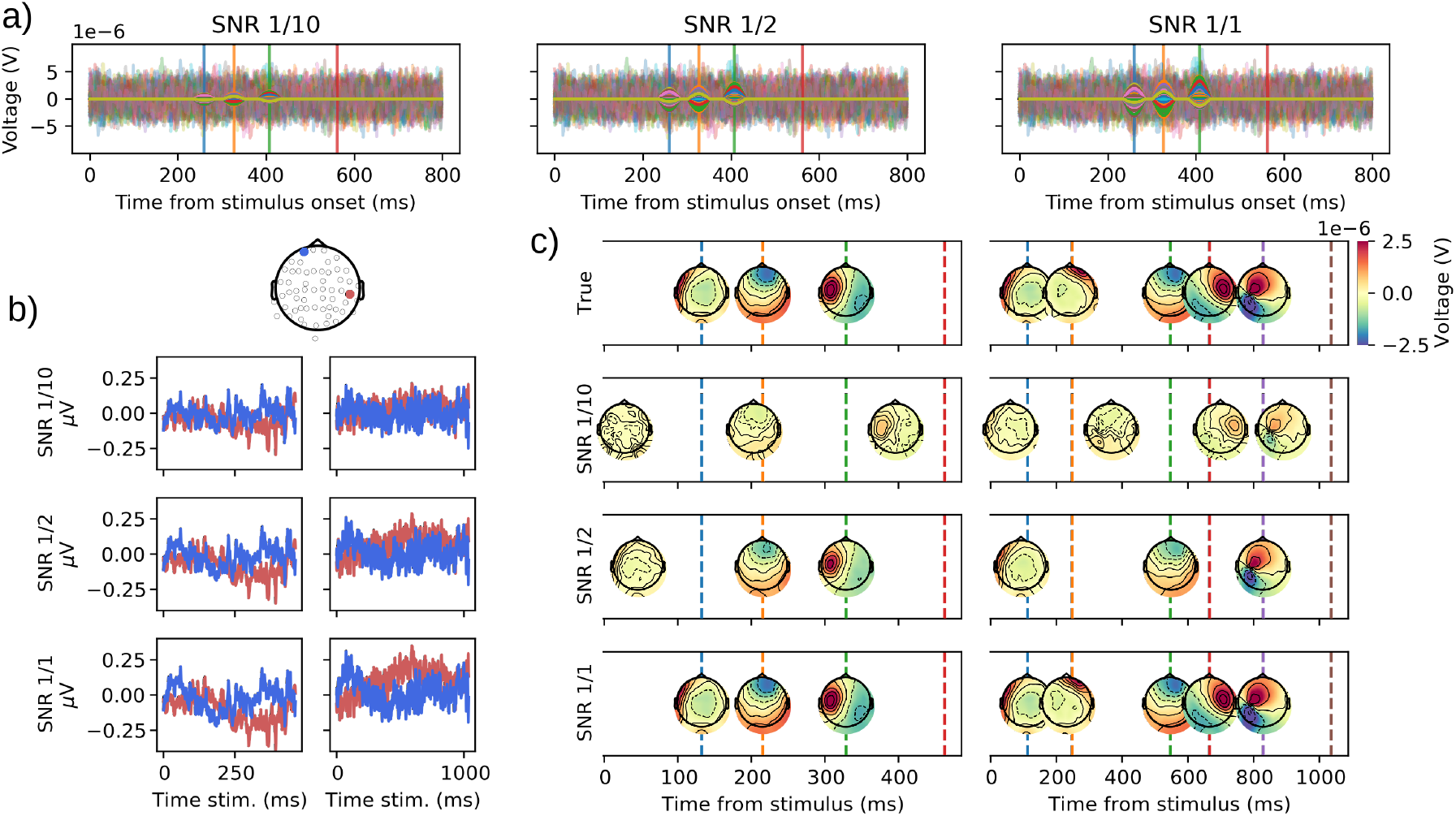
a) Illustration of the single-trial electrode activities yielded by three sources which peaks were simulated at the time points indicated by the vertical lines (red line indicates simulated behavioral response). The left-most plot shows the single-trial ERP for all electrodes of a three-event simulated dataset at an SNR of 0.1, where the single-trial ERP is displayed transparently and the underlying source activity without noise as solid lines. The middle panel represent a SNR of 0.5 while the right-most panel shows the single-trial event-related potential for a SNR of 1. b) Average ERP for the two electrodes (location depicted above the top row) for the three (left column) and five events (right column) simulated datasets for each SNR (increasing order). c) True (top row) and estimated HMP solutions for the datasets with three (left column) and five events (right column) and SNR of 1/10, 1/2 and 1/1.

All 6 datasets (3 or 5 events with SNRs of 1/10, 1/2 and 1/1) shared the same seed to illustrate how varying SNR and number of events changes the estimation while leaving stochastic elements unchanged between the datasets. The resulting simulated datasets were fitted with HMP using the default fitting routine described in Section 3.1.

Figure 3c illustrate the ground truth (top row) for 3 and 5 events compared to the estimation made with the three levels of SNR. As can be seen in the estimations, the recovery of the simulated sequence is almost perfect with an SNR of 1 (but see underestimation of the 2nd inter-event interval in the 5 event dataset). As expected, the match with the ground truth decreases with SNR. It is however noteworthy that even with a SNR of 1/10 and only 1000 trials, HMP still detects 3 and 4 events in the 3 and 5 events datasets even though the average times and topographies are fairly far from the ground truth.

It should be stressed that the impact of SNR is dependent on the number of trials (see for example recovery performances in Appendix A with only 100 trials). The higher the number of trials, the lower the SNR can be for HMP to perform well. HMP’s ability to identify events is also dependent on the ancillary assumptions of template duration and time distribution. To further explore the effect of SNR and its interaction with the number of to-be-estimated events, Appendix A parametrically explores the impact of SNR with different event numbers while appendices B and C explore the impact of different template duration and time distributions.

## 4 Application to real data

To test HMP on real data we analyzed three different EEG datasets. The two first datasets are extracted from the ERP-core database (Kappenman et al., 2021) and aim at showing how HMP relates to canonical ERP components such as the P3 and the N2pc. To show a less straightforward application than classical EEG paradigms, the last subsection is a dataset on a decision-making task from Boehm et al. (2014), previously analyzed by Van Maanen et al. (2021) using the original method by Anderson et al. (2016).

### 4.1 Method

#### 4.1.1 EEG recording and data preprocessing

The two datasets from the ERP-core database were measured within the same 40 participants using 28 electrodes (10/20 System). The datasets were already preprocessed, epoched, downsampled at 256Hz and high-pass filtered at 0.01Hz (see Kappenman et al., 2021, for specific preprocessing details). The only modifications compared to the original analysis is that the data was re-referenced to the average of the 28 electrodes, that no participants were excluded and no low-pass filtering was applied (outside of a 128Hz low-pass filter applied during the resampling procedure). The number of trials was 6179 and 9100, 168 and 228 on average per participant, respectively for the P3 and the N2pc datasets. A few trials (< 1%) were rejected as the response were given after the epoch length (> 800 ms).

The decision-making dataset was also already artifact-rejected and epoched. To conform with the study of Van Maanen et al. (2021) we applied an offline bandpass filter between 1 and 35Hz^1^ (one-pass, zero-phase, non-causal bandpass IIR filter, -6 dB cutoff frequency at 0.50 Hz and 39.38 Hz) using MNE (Gramfort et al., 2014). Noisy electrodes were marked as bad before the independent component analysis (ICA) and interpolated after ICA reconstruction. Electrodes were re-referenced to the average of all EEG electrodes. We then applied an ICA to repair eye-blink artifacts and used auto-reject (v0.4.2) to automatically interpolate epoch-wise bad channels or reject epochs (Jas et al., 2017). The data is composed of 30 electrodes (10/20 system) with 4547 trials for 25 participants, an average of 182 trials per participant. Most rejected trials were rejected based on autoreject analysis, a minor part of trials (< 1%) were rejected as the response was given after the epoch length (> 3000 ms).

#### 4.1.2 HMP estimation

For all datasets, the data was first truncated to the duration (RT) of each trial after adding a constant of 50 ms to the RT (adding samples after the RT allows to more easily detect event bordering the response). We performed a spatial PCA to the average variance-covariance matrix of all trials after centering the electrodes for each epoch. We then selected the first principal components that explained 99% of the variance (10 in both ERP-core datasets and 5 in the decision-making dataset) and zscored the principal components per trial.

The HMP model was then initialised as in Section 2.4: a pattern width of 50 ms, a gamma distribution with a shape of 2, similar to Van Maanen et al. (2021). The estimated parameters were the number of events, the scale of the gamma distributions and the multivariate contributions of each virtual channel for each event, equivalent to the simulation studies. The tolerance for the expectation-maximization (Equation 12) was set to 10^7^ to ensure most accurate estimations.

For all the conditions in all three data sets we fitted completely independent HMP models. For simplicity the HMP models were fitted on all the data without taking into account the participant dimension. For each analyzed dataset we followed the convention introduced in Figure 2 by displaying the average HMP time-course, the ERP centered on stimulus onset for a selected electrode (based on the contributions of each electrode to the relevant event), the ERP for that electrode centered around detected events and the distributions for event positions and between event intervals.

For all the HMP models estimated for the datasets, the times and topographical maps were computed based on the by-trial maximum probability for each event as in the simulations and as described in Section 2.2.4.

#### 4.1.3 Statistical analyses

To leverage the by-trial estimates, the condition effects in the N2pc and decision-making datasets were analyzed using linear mixed models. While these models might be more complex than the statistical methods used in most ERP research, they are the best way of exploiting single-trial measurement by accounting for the hierarchy in the data (participant level vs group level). The use of summary-based statistics (e.g. t-tests, ANOVA) is possible but they ignore the number of rejected trials per participant (thus all participant have the same weight irrespective of the number of trials retained) or different number of trials between conditions (e.g. the experimental design of typical P3 experiments). These linear mixed models were fitted in a Bayesian context, to avoid any convergence issues, using the bambi Python package (Capretto et al., 2022, version 0.13.0) with default priors weakly informed by the data, 4 MCMC chains, and 2000 samples. Posterior distribution are summarized by the point estimate of the maximum a posteriori and 95% credible interval (CrI) computed with the Arviz Python package (Kumar et al., 2019, version 0.17.0) based on the highest density interval (Kruschke, 2010). For simplicity we fit the models on the raw scales (*µ*V and milliseconds) and report the slope of the model as the difference between conditions. As the aim of the analysis is to briefly illustrate how HMP can be used in an experimental setting, we only reported the fixed effect of each linear mixed model. The full estimations files, convergence diagnostics and implementation details can be found in the associated OSF repository.

#### 4.1.4 Results

### 4.2 HMP and the P3

The P3 dataset from the ERP-core database was recorded from an adapted oddball task where participants had to indicate with a button press whether a letter was the target (Rare) or not (Frequent). In this paradigm the expectation is that a P3, an EEG-component believed to be involved in decision-making, is generated in the rare condition only. This component is a positivity typically observed at parietal sites in a window from 250 to 500 ms (Polich, 2007). We hypothesized that the Rare condition should yield an additional event with a timing and topography congruent with its counterpart observed in trial-averaged waveform.

To test this, we fitted two independent HMP models to the frequent and the rare conditions. Because the number of events is estimated from the data, this constitutes the test of the presence of an additional event.

As can be seen in Figure 4a, HMP finds 3 and 4 events respectively in the Frequent (top) and Rare (middle) condition. Therefore it appears that, as hypothesized, the Rare condition elicits a by-trial component not present in the Frequent condition. This additional event is a strong positivity over central/parietal electrodes (Figure 4a) with a much sharper waveform (Figure 4b bottom, third panel) than the stimulus centered ERP (Figure 4b top). This event occurs on average at 365ms after stimulus onset with a standard deviation across-trials of 77 ms (Figure 4c).

**Figure 4.**
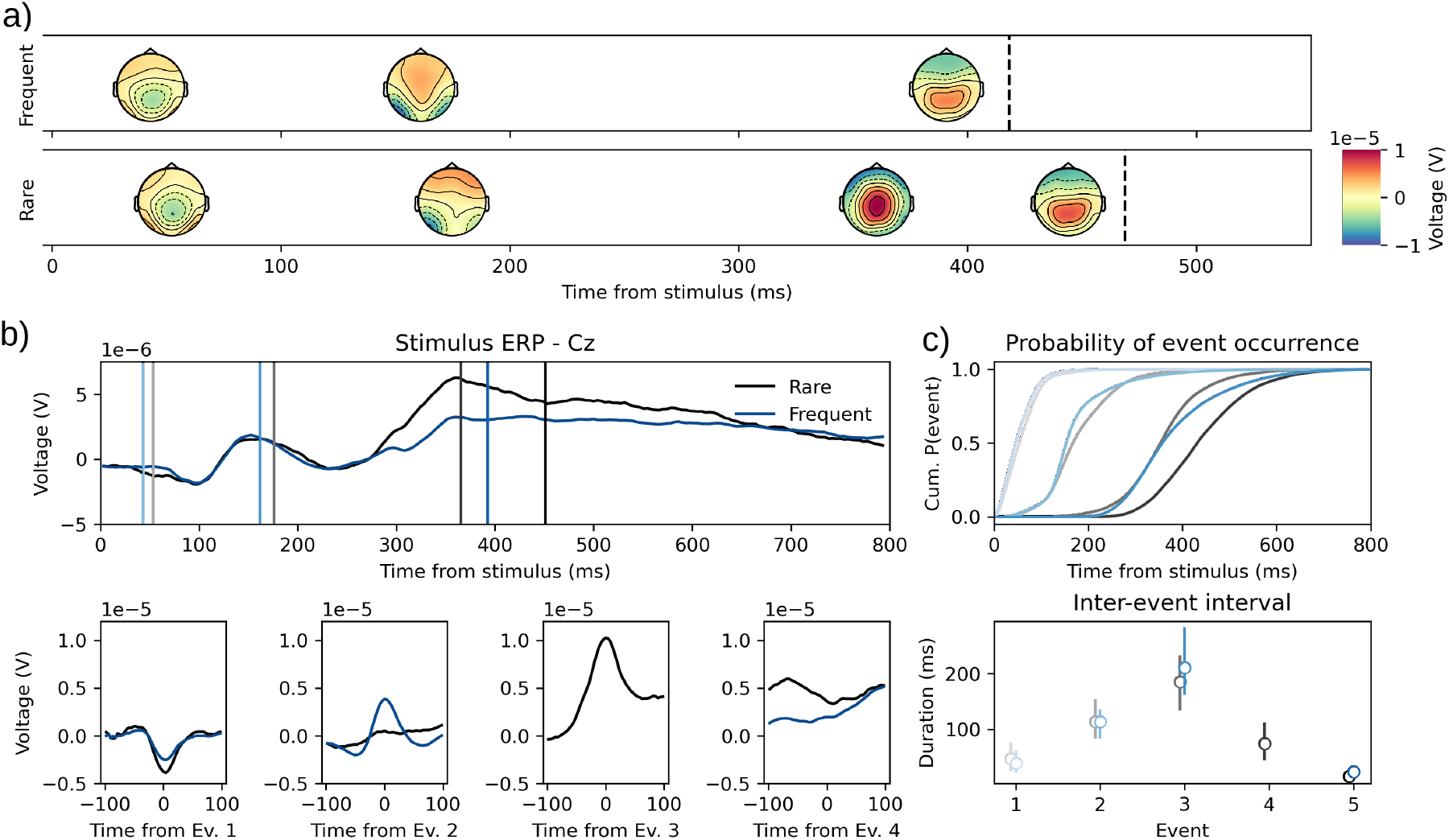
a) Average time-course and topographies for the identified HMP events in the frequent (top) and rare (bottom) stimulus conditions. Colored lines represent the average peak of the different events in each condition. b) Stimulus (top panel) vs HMP event (bottom panels) centered averaged ERP for the Cz electrode in both frequent and rare conditions. Ev.: Event. c) Occurrence probability in time from stimulus for the detected events (top) and distribution (1, 2nd and 3rd quartile) of the time between identified events (bottom). Black: Rare; Blue: Frequent; shading indicates sequence of events.

This P3 dataset analysis suggests that there is indeed an additional event in the rare stimulus condition, with a timing and a topography very close to the P3 usually observed on stimulus centered ERP. A notable difference of the peak-centered ERP is that the P3 is a much sharper component. Interestingly, the highest trial-averaged activity at each electrode suggest that this P3 component is located at the vertex (Cz) instead of more parietal electrodes such as Pz, which is the reported typical P3 location (but see the results in the decision-making dataset).

### 4.3 HMP and the N2pc

The N2pc dataset was recorded from a visual search task where participants had to indicate the orientation of a salient stimulus, presented either in the left or the right visual field among distractors. This paradigm is thought to elicit the N2pc, a component associated with attention orientation mechanisms (Luck & Hillyard, 1994). This N2pc has been found to be a difference in peak amplitude between contra- and ipsilateral sites with respect to the visual field where the target is located. Here, we hypothesized that this asymmetry would present in the topographies of the HMP identified events. We tested this using a linear mixed model on the peak voltage value for each event on the electrodes usually associated with the N2pc (PO7 and PO8, as in Kappenman et al., 2021). The electrodes were recorded as ipsi- or contra-lateral to the target side. The dependent variable was the difference in voltage for each trial between ipsi- and contraleteral sides. The model only included a fixed and random intercept term for each event. This statistical model therefore tests for the difference ipsi/contralateral in each event voltage while accounting for individual differences in these effects.

The results of the two fits on left vs right target presentation shows that very similar events are found between both conditions, in terms of topography (Figure 5a) and in terms of timing (Figure 5c). It appears quite clearly that the second event presents the expected asymmetry (Figure 5a, right panel). The maximal of the by-trial averaged signal amplitude for this second event is located on electrode PO7 and PO8 respectively for the left and right field HMP model. When recoding the electrodes as contra- or ipsilateral to the attended visual field and computing the differences for each event it appears that the ipsilateral side has a higher voltage specifically on the 2nd event (2.53*µ*V, CrI = [2.03, 3.00], Figure 5b). When inspecting the other events it appears that the first event also presents a difference but with a (slightly) more negative voltage for the ipsilateral side (−0.32*µ*V, CrI = [-0.57, -0.07]). The two last events, however, do not display evidence for a difference between ipsi- and contra-lateral electrodes (3rd event: -0.50*µ*V, CrI = [-1, 0.02], 4th event: 0.09*µ*V, CrI = [-0.33, 0.47]). Thus the N2pc dataset generates only one HMP event that displays the expected visual field asymmetry and further suggests that the event prior to the N2pc also displays an asymmetry in the opposite direction. If generalized to other datasets this early effect could reflect the fact that early visual potentials are already sensitive to the hemifield of the target, at least when it is displayed with a different color as in the dataset by Kappenman et al. (2021).

**Figure 5.**
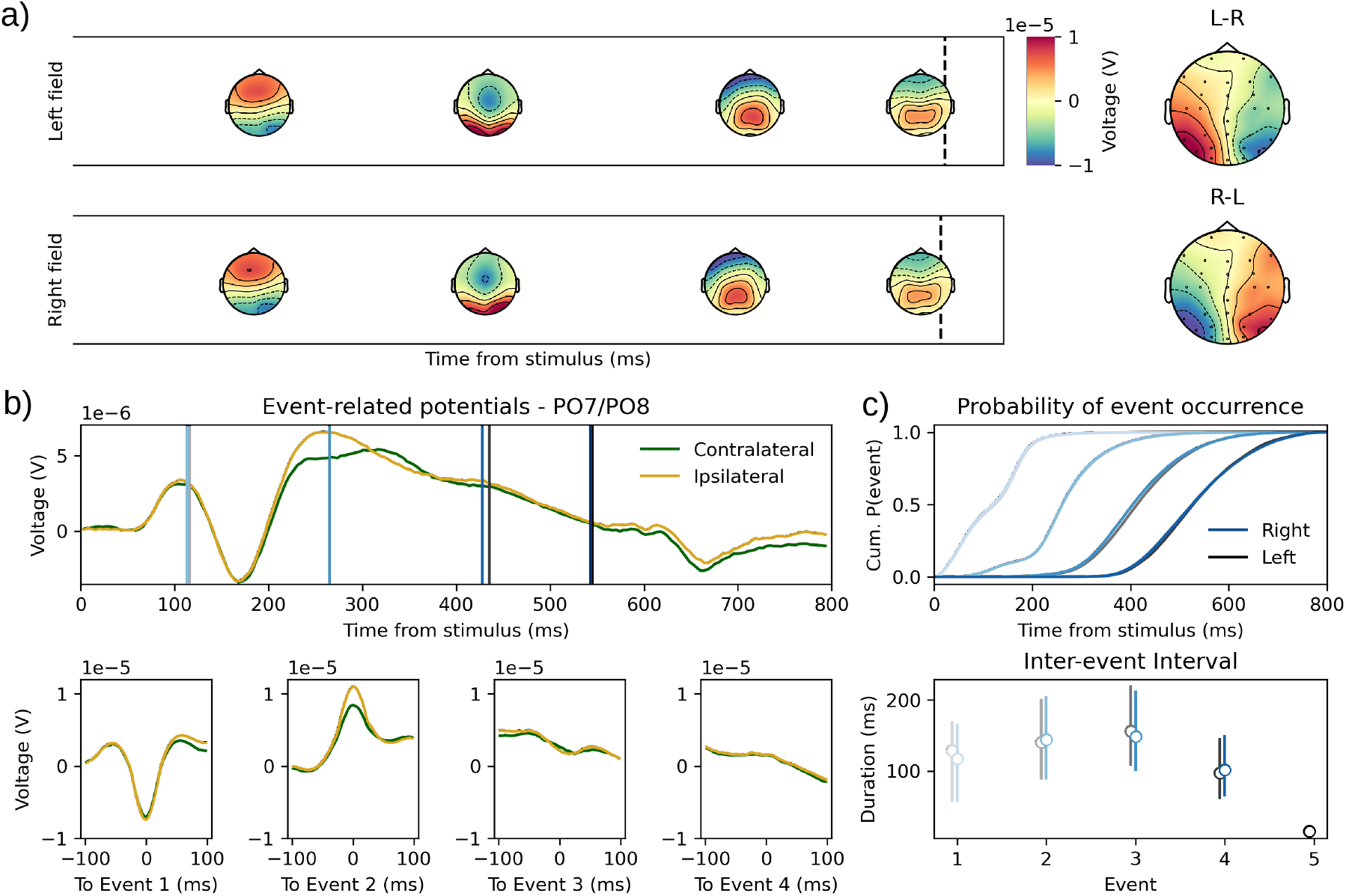
a) Average time-course and topographies for the identified HMP events when the target was presented in the left field (top) vs the right field (bottom). The right most plot shows the difference in topographies of the second event when subtracting right field to left field (top) and vice-versa (bottom). b) Stimulus (top panel) vs HMP event (bottom panels) centered averaged ERP. The electrodes were recoded to contra- and ipsilateral depending on the side of the target. The blue and black lines represent respectively the average times from the events detected in the right and left field models. c) Occurrence probability in time from stimulus for the detected events (top) and distribution (1, 2nd and 3rd quartiles) of the time between identified events (bottom) for the left and right visual field presentation. Black: Left; Blue: Right; shading indicates sequence of events.

### 4.4 HMP and decision-making

#### 4.4.1 Introduction and methodology

In this decision-making task, participants were asked to indicate whether the majority of points in a cloud of moving dots (random dot kinematogram) was moving to the left or to right with the corresponding hand. The speed accuracy tradeoff was manipulated with a cue indicating to the participant, at each trial, to either favor the speed or the accuracy of their response (see Boehm et al., 2014, for further details on the experimental protocol).

Using a LOOCV procedure on different fits of the method of Anderson et al. (2016) with different assumed number of events, Van Maanen et al. (2021) found that a two-event model fitted the speed instruction data best while a three events model fitted the accuracy instructions best. The authors interpreted this as evidence that an additional psychological process was present in the accuracy condition. Van Maanen et al. (2021) also found that only one between-event interval was sensitive to the speed instructions and therefore interpreted that this interval was the period during which the decision was taken. The original study used the equivalent HMP parametrization described in Section 2.4 on 100 Hz downsampled EEG data. Here, we attempted to replicate their findings using HMP with the same parametrization but using the sampling data at which the EEG was acquired (500 Hz)^2^ and inferring the number of events from an HMP fit as described in Section 2.3.1.

In this dataset the nature of the events to be found is less straightforward than the two previous applications. We therefore used the estimation made of the N2pc and the P3 in the previous datasets to test whether events close to these components are found in the new dataset. For this, we computed the difference between the averaged topographies estimated for the identified N2 (the second event of an HMP model of section 4.3 for both left and right field combined) and P3 (the third event for the HMP model estimated for the rare condition in Section 4.2) with the averaged topographies of all events in the estimated HMP models for speed and accuracy conditions. These differences were computed for all common electrodes between both datasets (N = 17). We then classified each newly discovered event in both conditions of this dataset as being similar to a N2, a P3 or unidentified (if the event was closer to a 0 activity virtual event or less close to the components than other events in the same condition) based on the mean squared deviation with the N2 and the P3 topographies.

In addition to an eventual N2 and P3, given that responses were provided by participant with their right and left hands, we expected that an event related to response execution would be present in the HMP solutions. Given usual observations on lateralized readiness potentials, we expected that the response preparation event would display a negative correlation between left and right hemisphere electrodes on the voltage difference between left and right responses. To compute this correlation we first computed the difference for each electrode between left and right responses for each averaged event peak. The correlation was then computed between the left hemisphere and right hemisphere electrodes (excluding all midline electrodes) for each event in each speed-accuracy conditions. Statistical inference was done on Pearson’s correlation coefficient, where the alternative hypotheses is that the correlation coefficient is negative with a statistical threshold of *p* = 0.05. All described classification attempts could not be done per trial and instead have been done by averaging simultaneously over trials and participants for simplicity, thus effectively treating all trials in the dataset for each condition as stemming from the same distribution.

After event classification, we tested the impact of speed-accuracy instructions on the by-trial time between each event. To draw statistical inferences from the time difference between event peaks, we fitted a linear mixed model, as described in Section 4.1, on the intervals in milliseconds between events without intercept and with factors event number (categorical) and their interaction with speed instructions as predictors.

#### 4.4.2 Results and discussion

As can be seen in Figure 6a, HMP detected 3 events in the speed condition and 4 events in the accuracy condition. Computing the mean squared distance between the topographies of the detected events and the topographies of the N2 and P3 from the previous sections, showed that the shared 2nd event in both conditions and the 3rd event in accuracy-focused condition are most similar respectively to the N2 (Figure 5a) and the P3 (Figure 4a). Noticeable differences with the previous P3 is that 1) the timing is slower for the new P3 with a mean time of 518ms (SD = 193ms) after stimulus onset and 2) that the positivity is more parietal with the maximal activity at electrode Pz. Regarding electrode asymmetry (Figure 6a right-most panel) with respect to response hand as evaluated by a Pearson correlation coefficient, the last event in the accuracy condition (*r*(11) = −0.60, *p* = 0.02) as well as the last event in the speed-focused solutions (*r*(11) = −0.98, *p <* 0.001) display the expected negative correlation between hemispheres but none of the other events (all *r* ≥ 0.87).

**Figure 6.**
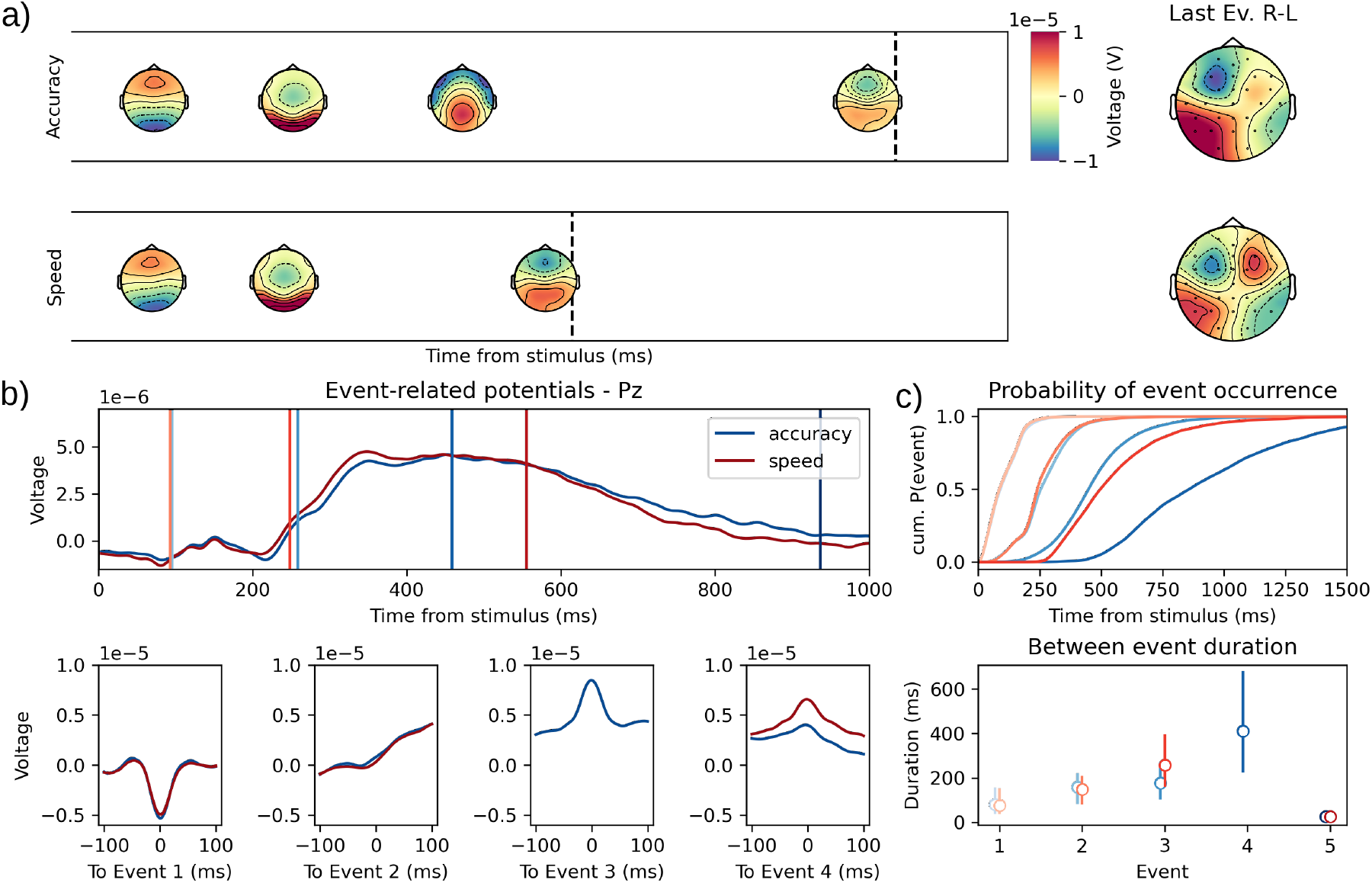
a) Representation of the HMP solutions for Speed (top) and Accuracy (bottom) conditions. The right-most panels represent the difference in topographies for right vs left responses for the HMP event estimated before the response (color of the contributions for these two topographies are scaled between -1 and 1 µV). b) Average ERP on the Pz electrode centered on stimulus onset (top) vs the HMP detected event (bottom). Colored lines represent the average peak of the different events in each condition. c) Occurrence probability in time from stimulus for the detected events (top) and distribution (1, 2nd and 3rd quartiles) of the time between identified events (bottom) for the speed and accuracy instruction conditions. Ev.: Event; Red: Speed; Blue: Accuracy; shading indicates sequence of events.

To analyze the difference in time (illustrated in Figure 6c) for each event between the speed-accuracy stress conditions, we ignored the additional HMP event in the accuracy condition as it does not have an equivalent in the speed condition. Given that the main manipulation of the task is on the speed stress imposed to the decision of the participants, we analyzed the time differences between all successive events to locate where in the detected RT intervals do the participant speed up. From the analysis of the by-trial intervals between events it appears that no evidence is found for an effect of speed instructions on the first time interval, from stimulus onset to the first event (*β*_1_ = -2.69, CrI = [-11.55, 5.16]), nor the last time interval from the last HMP event to the response (*β*_4_ = 0.25, CrI = [-7.27, 7.99]). The second time interval, from the first to the identified N2, is shorter in the Speed condition but the evidence is low as credible intervals loosely contain 0 (*β*_2_ = -7.99, CrI = [-17.76, 2.18]). Finally, the time interval from the before last event (N2 in speed, P3 in accuracy) to the last event, is highly changed by the speed instructions (*β* = -168.61, CrI = [-223.62, -113.98]).

To further showcase how the method can be used in an exploratory setting, we choose to analyze the single-trial peak voltage value at the Pz electrode (Figure 6b), as it maximal for the additional event in accuracy, using the same linear mixed model as for the inter-trial intervals. Doing so we see that the first event already displays a less negative peak although the difference *µ*V is quite small (*β* = 0.42, CrI = [0.04, 0.78]). The N2 event could be slightly more negative but the credible intervals loosely include 0 (*β* = -0.52, CrI = [-1.09, 0.01]). The biggest difference however, is observed for the last event before the response where the voltage is much higher in the speed condition (*β* = 2.53, CrI = [1.89, 3.20]).

To sum up, we identified one additional HMP event when participants were required to be as accurate as possible. Thanks to the two previous datasets we could categorize this additional event as a P3. The reason why this P3 component is not found in the speed-focused condition is not the scope of the current manuscript, nevertheless, it is worth noting that this result is coherent with a study by Kutas et al. (1977) showing that a P3 is not found before the response in a significant proportion of trials exclusively in the speeded RT condition. The authors argued that the extreme speed stress would make the P3 and response production appear simultaneously which would be congruent with the increased peak in Pz at the moment of the response-lateralized event. A last note is that while we replicate most of the results of Van Maanen et al. (2021), we do observe an additional event that we could relate to response production, thus showing that HMP is more sensitive than the method used in the original study.

## 5 Discussion

In this paper, we introduced HMP, a method intended to detect hidden single-trial events in neural time-series. The method uncovers these events by assuming that the data is made of sequential multivariate patterns, repeated across trials with varying timing respective to the previous, external (e.g. stimulus) or hidden, event. The number of events, the per-event multivariate representation on channels and time distributions are estimated from the data. We believe that this approach is a significant step towards resolving distortions related to averaging in neural time-series (see Luck, 2005; Mouraux & Iannetti, 2008, and Section 3.2). After describing the method, we showed in simulations (see Section 3 the core assumptions of HMP and the consequences of their violation (see Section 3.2), and then showed the method’s ability to recover by-trial EEG events under various levels of signal-to-noise ratio even with only 1000 or 100 trials (see respectively Section 3.3 and Appendix A). In a last section we applied the method to three previously published datasets, two datasets from the ERP-core database (Kappenman et al., 2021) dedicated to specific ERP components (P3 and N2pc), and one dataset with less straightforward expectations regarding the nature of the events present in the data.

The analysis on the P3 dataset in Section 4.2 showed that HMP detects an additional event when a stimulus is rare compared to when a stimulus is frequent. The HMP analysis further allows to characterize this additional event as corresponding to the P3, based on its by-trial peak voltage and its by-trial time location. For the N2pc dataset in Section 4.3, HMP showed that the single-trial electrode activity around the 2nd detected event presents an hemispheric asymmetry depending on the hemifield in which the target was presented. Finally using HMP we could locate these two components and a response-lateralized component in a more complex decision-making task in Section 4.4. The interval between the N2/P3 and the response-lateralized event contained most, if not all, of the effect of speed-accuracy instructions. Overall this last section shows the potential of HMP not only from a signal processing perspective but also from a psychological perspective as, assuming that N2pc and P3 reflect respectively attentional filtering (Eimer, 1996) and decision-making (Polich, 2007), we then have single-trial estimates for the time at which these processes are present in the RT.

Importantly all statistical analyses performed on these HMP events did not necessitate to rely on arbitrary rules on the averaged ERP waveform to conclude on an effect of interest (e.g. a *µ*V threshold, J. Miller et al., 1998). In fact no information from the averaged ERP was used to derive HMP events, only the assumption that single-trial events are sequential multivariate patterns allowed to reach similar conclusions than well-known ERPs components. With these assumptions, the peak of any HMP event is given by-trial and can be readily used to test hypotheses such as the timing of an event or the electrophysiological profile, as shown throughout the analyzed real datasets.

### 5.1 Mental chronometry considerations

The obvious target of HMP estimations are questions related to mental chronometry, the study of the timing of mental processes. As an example, using a previous iteration of the method, Van Maanen et al. (2021) successfully fitted an evidence accumulation model to the estimated by-trial times the events associated to the decision. The range of questions is quite large as one can then use the interval between detected events and test for experimental effects based on expectation for attention filtering, motor preparation and so on. Note that contrary to most classical methods from mental chronometry (Brown & Heathcote, 2005; Donders, 1868; Ratcliff, 1978; Stone, 1960), HMP does not imply seriality in cognitive processes. This means that a particular process is not expected to end at the start of the following event. Instead, the time between two events at best predicts the time needed for the following cognitive process to start. In other words, common assumptions about parallel processing or coactivation (McClelland, 1979; Miller, 1993; Townsend & Nozawa, 1995) are perfectly compatible with HMP as long as the peaks of the events follow each other. For example in decision making, the minimal assumption is that a decision follows sensory encoding of the stimulus and precedes the response execution. There is no strict need to assume that a decision process cannot continue during response execution, compatible with observations from a growing number of studies (Buc Calderon et al., 2017; Selen et al., 2012; Servant et al., 2021; Thura et al., 2022; Weindel et al., 2022).

### 5.2 Additional signal processing methods facilitated by HMP

The assumption and development of HMP has been driven by work on mental chronometry and cognitive modelling but it is important to stress that the benefits of HMP go beyond these application domains. Indeed, many applications in cognitive science suffer from jitter in the time of the underlying cognitive-related events in EEG, MEG, and other related measures. By explicitly making the assumption of sequential events in the brain following estimated probability distributions, HMP is able to recover the underlying sequence. This allows researchers to center the signal around the event of interest and perform any type of analysis, whether it is about connectivity (Portoles et al., 2022), inverse modelling, time or voltage analyses as in the present example application, spectral composition, or otherwise. As another example, with these by-trial peaks it is also possible to use single-trial predictors (e.g. a wide range of continuous stimulus intensity) to estimate the effect of that manipulation.

Apart from the timing information of events which can inform a lot of applications in neurophysiological signal analyses, it should also be noted that the spatial information on the channel contribution to each event can also be leveraged for further processing. The topographies reported for the HMP estimation in the present paper naturally lend themselves as spatial filters, turning complex multivariate analyses to 1-dimensional timeseries or time-frequency series.

### 5.3 When to apply HMP

The question of the application of HMP is then reduced, as illustrated by Section 3.1, to its core assumptions which can be summarized as follows: is the sequential nature of these transient and trial repeated events reasonable given the task of the participants? If so, and if the data have a proper signal-to-noise ratio similar, if not lower, to what one would require for traditional ERPs (see Section 3.3 and Appendix A), HMP will work well. Of course, the ancillary assumptions of probability distribution and pattern width expectation, explored in Appendices B and C, can have an impact on the successful identification of the cognitive events. In particular, violation of the expected pattern duration (Appendix B) can have consequences: If (and only if) the actual underlying pattern is shorter than the one that is expected, the true positive rate decreases, meaning that not all events are identified. We believe that the initial duration of 50 ms chosen by Anderson et al. (2016) is reasonable given the other studies that successfully applied it. Nevertheless, the width of a pattern can change to quite some extent depending on the cells and neurons actually recorded (e.g. MEG vs EEG vs intracranial electrodes). This pattern can also change depending on the stimuli used, for example spatial frequency of visual gratings have been shown to have a broad impact on scalp topographies as seen using EEG (Kenemans et al., 2000). This latter point invites caution when assuming a pattern width even within a given measure for which the usual pattern width works well.

To be sure, running HMP with the minimum possible duration (but defined on enough samples to reliably estimate the pattern) will prevent missing events in the data. In this case, HMP will likely duplicate events that are longer than the expected pattern but researchers can either progressively increase the width of the searched pattern or rely on alternative, second-step, procedures such as LOOCV (see for example Borst & Anderson, 2024). Such a latter approach would reveal whether the model identifies more events than are present in the data.

The method can be generalized to any template that is expected in the time-series. If, for example, certain applications are expected to generate biphasic neuronal responses, the equations in Section 2.4 can be adapted to reflect this expectation.

On the distributions of the times between the events, as shown by Appendix C the choice of a specific distribution does not dramatically impact HMP performance. Overall it is important to note that the chosen distribution helps in the identification of the event, but the resulting by-trial estimation of times will only resemble the chosen distribution’s shape if there is not a strong signal-to-noise ratio (e.g. if true underlying distribution is left-skewed but a target distribution with a right-skew is used, providing enough signal, the estimated by-trial event distributions will resemble a left-skewed distribution).

A source of limitation in HMP is the minimum time imposed to the average interval between events described in Section 2.2.1. This minimum average difference in time between events is a limiting factor to detect events very close in time (e.g. if the interval between two successive events is on average faster than 50ms). This is illustrated by the impact of the average RT on the performance of HMP in Appendix C. This limitation can however also be alleviated by either searching for shorter patterns or reducing this constraint (e.g. signal-to-noise simulations in Appendix A is even better if this censoring is reduced for half a pattern width) and then apply second step procedures to filter out duplicated events (e.g. re-fitting an HMP model with only the event displaying a low correlation between them).

A final word on the conditions to use HMP is the measure of the end of the sequence. In this manuscript we always used the time of the response as the signature of the end of the sequence of events. It would however be perfectly possible to use other sequence end times externally measured (e.g., electromyography, oculometry,…) as long as the chosen index is informative about the end of the sequence and of high temporal resolution. If an index is chosen without information about the end of the sequence of interest (e.g. the start of the next stimulus), it is likely that the estimation of HMP will not perform optimally (e.g., Figure 2b). Hence, a well defined by-trial time for the end of the sequence is a necessity to apply HMP.

### 5.4 Comparison to other event identification methods

Discovering recurring patterns in the EEG is not a new idea. Lehmann (1971) showed how EEG time-series can be summarized in a small number of so-called micro-states, i.e. quasi-stable states of electrical potential across electrodes, typically 80-150 ms (Brunet et al., 2011; Koenig et al., 2002; Michel & Koenig, 2018; Van de Ville et al., 2010). Each microstate is therefore defined by its EEG scalp topography. In the past, four canonical topographies have been identified, under which most microstates can be classified (Michel & Koenig, 2018). Microstate topographies and their distributions are usually identified based on group-averaged data, after which the samples in the original EEG are labeled accordingly.

This highlights the main difference with HMP, which identifies events on a by-trial basis, and then averages the results (e.g., Figure 6). This is crucial, as averaged data, particularly of longer trials and samples further away from fixed time points, might obscure important events (Borst & Anderson, 2024; Luck, 2005). This is, for example, clearly visible in Figure 2c, where distinct trial-by-trial events result in very weak averaged ERPs. We hypothesize that the events that HMP detects might be the peaks of the microstates found in averaged data (which is the reason they are quasi-stable for some time). Although we think that the by-trial estimation of HMP is an advantage for task-based data, so far HMP cannot be used to analyze resting-state data – an area in which the microstate analysis has been particularly successful (Michel & Koenig, 2018).

Another commonly used method that can be linked to the results of HMP is hidden Markov modeling (HMM). HMMs start to be widely applied in the fields of neuroscience including on EEG and related measures (Kirchherr et al., 2023; Masaracchia et al., 2023; Quinn et al., 2018). Contrary to HMP, HMMs assume a fixed number of states and generative distributions (e.g. for the electrodes) specific for the states. Despite their resemblance, HMP is not a specification of a HMM. Their common ground is that both can be described as transitions between states. However, in HMP these transitions are signaled by short-lived multi-variate patterns and can only happen from one state to the following one. Additionally, HMMs can transition to any state but assume that the state’s data generating distribution is present during the whole activation of the state.

Although HMMs can be very useful in the context of cognitive neuroscience, their difficult estimation and parametrization can lead to volatility in the estimation and quick erroneous switches between states (Masaracchia et al., 2023). For this reason, we believe that HMP, as long as the assumption of sequential events is met, provides an easier and more robust estimation of a cognitive event sequence in the context of the timing of these events. As a final note, given a correct specification of both HMM and HMP and if all underlying events are sequential to another, both methods should converge to the same solution.

Another approach is to regress out EEG activity based on external events, such an approach has been implemented in the unfold toolbox (Ehinger & Dimigen, 2019). Using this toolbox Froemer et al. (2022) have shown how common trial-averaged stimulus centered ERPs can show misleading activity as to their relation with behavior. This method’s objective is to deconvolve components from the EEG using linear deconvolution modeling based on external events (e.g. stimulus or response onset). As can be seen in the noiseless simulation of Section 3.1, the overlapping of components can be the consequence of time-jitter in the underlying single-trial events. Therefore this method could be combined with the internal events detected by HMP to further improve the characterization of the average EEG response.

Other methods have been used to specifically obtain single-trial measurement of EEG or MEG components to achieve RT decomposition. For example, Ruchkin and Glaser (1978, as cited by Smulders et al., 1994) looked for a pattern of a half-sine with a duration varying from 250 to 970 ms to achieve P3 single-trial measurements. For the N2 also single-trial measurement has been proposed by Nunez et al. (2019) this time based on a singular value decomposition of the average ERP. In the case of the detection of a single specific event, such single components methods could outperform HMP as they benefit from looking at known exact properties from a specific component.

### 5.5 Implementation

A final note on the implementation of HMP is in order. We created an open-source and collaborative Python package (https://github.com/GWeindel/hmp). All results and plots presented in this study are based on this package (v.0.4.0). The repository contains tutorials on the data format, the fit of HMP models and the comparison of experimental conditions. This paper and the online repository should make the method more accessible for a broad range of interested researchers. In terms of computational complexity, fitting an HMP model can be quite expensive in terms of processing time and RAM. Dimension reduction through PCA and resampling have no dire consequences as long as the range of data reduction is in line with the desired accuracy.

In terms of the number of trials required to fit an HMP model, a precise definition lacks. The optimal number of trials will vary depending on the signal to noise ratio, the number of events expected and therefore the length of the RT (as illustrated in the signal-to-noise simulation in Section 3.3 and Appendix A) and the coherence of the sequence across trials. Despite these complex issues to define an optimal number of trials (e.g. as illustrated in the varying proportion of trials with a given event in Section 3, we can nevertheless provide some guidelines. The experimental data analyzed in the last result section with HMP is based on approximately 2200 trials per condition (both conditions are fitted independently). Per participant, this amounts to around 90 trials, which is usually on the lower end of trial numbers for many standard experiments in cognitive neuroscience. Based on this example and the simulations it appears that HMP does not require an excessive amount of trials. Moreover, the EM algorithm used also allows to easily fit conditions or participants with lower number of trials. Indeed, in the case of individual participants for example, by providing starting points that are estimated on the group level, it is possible to get reliable estimates even with a low amount of trials. This could be very useful for clinical settings where only a few data points per participant are available but enough participants have been recorded and analyzed to allow for a restricted search in the parameter space.

Although the current paper frames HMP as a method for EEG time-series analysis, analysis of other neural time-series data is also possible. The adaptation to MEG is already implemented within the Python package associated with HMP. Other time and chronometrically honored measures such as intracranial EEG and electro-corticography require, to date, a very small set of adaptations. The main challenge in these applications is the expectations regarding the pattern. In all of those applications, a half-sine seems as justified as for EEG but the duration of the expected event requires some adaptation due to the differences in sensitivity to different type of neural signals and to skin and skull distortions. Other time-series such as electromyography or pupillometry could also be used in an HMP model, but the relative high signal-to-noise ratio and the limited number of channels usually associated with those techniques, render the HMP method probably less useful than existing methods specifically designed for such measures (e.g. https://github.com/lspieser/myonset for electromyography or Cai et al., 2023, for pupilometry).

### 5.6 Future work

We plan to increase the flexibility of HMP to allow researchers to test different hypotheses and use HMP in different scenarios. For example, a parametrization with several expected event widths or probability distributions would allow one to progressively build task models that would capture complex sequences of events. Ultimately, these could be used as priors for HMP when analyzing similar tasks. This could even be taken to another level by directly integrating cognitive models such as ACT-R or evidence accumulation models within the estimation procedure (Turner et al., 2017). Another addition that is currently being studied is the possibility to fit a hierarchical version of the model where the estimation of group parameters inform individuals and viceversa (Lee & Wagenmakers, 2014). This would allow a cumulative approach to HMP applications by using priors and population distributions to constrain individual estimations. Lastly, for now HMP requires an RT for each trial in order to extract the event sequence, but our intention is to allow the method to accommodate different scenarios without requiring a response from a subject. This includes passive stimulus exposure, decisions without overt actions, or resting state data. This would open up the possibility to further the knowledge gained from classical mental chronometry to new applications.

### 5.7 Conclusion

By building on the method developed by Anderson et al. (2016) we show how HMP fitted on time-based neural signals can accommodate many different scenarios and robustly determine a sequence of neural events. The sequential nature as well as the use of probability distributions to enhance portion of the signal gives a high granularity in the interpretation of a series of stimulus and response related events in the brain. Given its qualities, this method represents a new venue for fundamental and applied research relying on reaction time based tasks, not solely for mental chronometry related questions, but also for a wide variety of brain-related signal analysis as advocated by the modern mental chronometry (Meyer et al., 1988).

## Data and Code Availability

All the data and code used to perform the analysis and generate the figures can be found at https://osf.io/29tgr/.

## Author Contributions

GW, LvM and JPB conceptualized the method. GW and JPB implemented the method in the associated python package. GW wrote the first draft of the manuscript. LvM and JPB reviewed and edited the manuscript.

## Acknowledgments

We thank John Anderson, Michael Nunez and Joshua Krause for the discussions about the method and beyond. We also wish to thank Leon Kenemans for providing insightful comments on a first draft of this manuscript. The authors wish to thank the COBRA and CogMod groups, respectively from Utrecht University and Groningen University, for early feedback on the development of the method. The authors would also like to thank the anonymous reviewers whose comments greatly improved the original manuscript.

## Appendix A

### Parametric manipulation of signal to noise ratio

To further investigate the effects and interactions between different SNR and numbers of to-be-estimated events, we simulated 500 EEG datasets. One dataset consists of 100 trials across 59 EEG channels. The events were defined as a 50 ms half-sine activation of the selected source and the by-trial between event intervals where all sampled from a gamma distribution with a shape of 2 and a scale of 100 (hence an average interval of 200 ms).

We manipulated the SNR of the events by drawing a source activation from a log-uniform distribution between 10^−8^ and 10^−6^ ampère for each of the 500 simulated datasets. The signal and amplitude at the electrode level varied depending on the sources location in the brain. The SNR was computed as described in Section 3.3. An average SNR was then computed per dataset based on the SNRs of all the simulated events within that dataset.

### SNR Simulation sensitivity measures and statistical analysis

To understand under what circumstances HMP can successfully recover events we first fit an HMP model with a given target pattern width and an expected distribution shape, to each simulated dataset using the cumulative event fit described in Section 2.3. The only difference is, to avoid too many false positives in the signal with very low SNR given that the local maximum approach for a 1-event model will nearly always find a local maximimum to partition the time-series^3^, we introduce a statistical test where the log-likelihood value of the first event is compared to the log-likelihood distribution of 100 random starting without the maximization step. The first event is retained if the log-likelihood is larger than 1.96 standard deviation of this distribution (*p <* 0.05). The events estimated this way are then compared to all simulated topographies by computing Euclidian distance between the activity in the channels (here principal components) at the true peak times and at the estimated peak times. We then categorized the events that are closest to a generating event by computing the median absolute difference with each event. This separated estimated event as successfully recovered events (i.e., hits), and false alarms (e.g. with a higher distance to generating ones or closer to a null channel event). From this we can express the success of the recovery for each simulation by the true positive rate (TPR, *i.e*., ratio of correctly identified events over total number of simulated events) and the positive predictive value (PPV, *i.e*., ratio of correctly identified event over the total number of identified events). Figure A1a illustrates two hypothetical examples with either a lower TPR (middle panel) or a lower PPV (bottom panel) compared to the true model (top panel).

**Figure A1.**
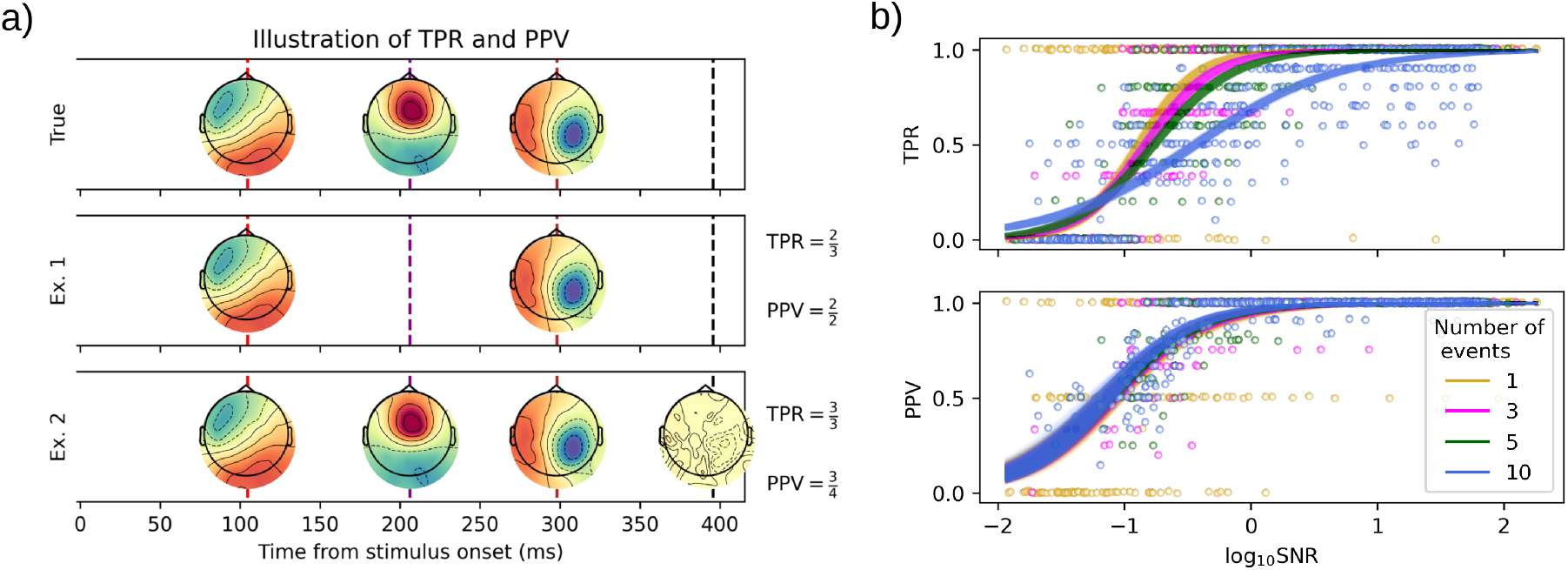
a) Two examples of estimated HMP models based on a simulated model (top). For the purpose of illustration, these estimations were done by forcing the number of events to be lower or higher than the actual number of simulated events to show the example of a missed event and an additional event detected by HMP. For Example (Ex.) 1, computing the electrode activity distances between the simulated events and the recovered ones suggest two correctly recovered events hence a True Positive Rate (TPR) of 0.66 but a Positive Predictive Value (PPV) of 1 (as all found events have a corresponding true event). In Example 2, this same computation outputs a TPR of 1 (as all events to be found were found) but a PPV of 0.75 as one of the events found doesn’t match any simulated event (or less so than the other). b) Results on the TPR (top row) and PPV (bottom row), the different number of simulated events are indicated by colors, fitted lines represent 1000 regression lines drawn from the posterior distribution of the fitted binomial regression.

The effect of the SNR was assessed by fitting a binomial logistic regression models to TPR and PPV. All coefficients of these regression models are expressed on the logit scale. Posterior distributions are summarized by the point estimate of the maximum a posteriori and 95% credible interval (CrI) computed based on the highest density interval (Kruschke, 2010). All regressions were fitted using the bambi (Capretto et al., 2022, version 0.13.0) Python package with default priors weakly informed by the data, 4 MCMC chains, and 2000 samples. The graphical summary of the regression model coefficients was done through the Arviz Python package (Kumar et al., 2019, version 0.17.0). Posterior distributions are summarized by the point estimate of the maximum a posteriori and 95% credible interval (CrI) computed based on the highest density interval (Kruschke, 2010).

### Results

For each of the SNR level we simulated a sequence of 1, 3, 5 or 10 HMP events (and a response production event). Figure A1b shows the TPR and PPV that we obtained for each of these combinations, with two binomial regression models to illustrate the trend. Both regression models take SNR and number of events and their interaction as predictors with a logit link, and TPR and PPV, respectively, as dependent variables.

Given the coding applied to the predictors, the intercept of the binomial regression is the expected TPR or PPV, on the logit scale, for datasets with 3 simulated events and for an SNR of 1. Slopes can be interpreted as the difference when increasing one log 10 unit in SNR or adding one event in the generated dataset.

For the TPR, the value of the intercept is equal to 3.14 (95% CrI = 2.78, 3.28) on the logit scale, meaning a TPR of 95.6% for a 3-event model at a SNR of 1. The slope of the SNR is equal to 3.79 (95% CrI = 3.53, 4.06) which means that the predicted TPR at a SNR of 10 is higher than 99.99% while a SNR of 0.1 (left panel of Figure 3a) is around 34%. The number of events decreases the TPR (−0.34, CrI= [-0.38, -0.30]) but interacts with the amplitude of the signal (−0.29, CrI= [-0.33, -0.24]), meaning that to obtain a good TPR, a higher signal amplitude is required for a higher number of events.

With respect to the PPV, the value of the intercept is equal to 3.02 (CrI= [2.78, 3.28]), meaning a PPV of 95%. All probability mass of the posterior distributions for the slopes are positive, meaning that increasing the number of events, or the signal’s SNR increases the PPV (see OSF repository for the full table of coefficients).

To sum-up, this simulation showed that HMP is able to recover the ground truth of a sequence of events, with few false-alarms, even in a time-series with very low signal-to-noise ratio and only 100 trials. To further test the impact of different parametric variations on the recovery of an HMP model, Appendices B & C respectively vary the deviation of the target pattern and deviation from the assumed time distribution family with a noisy signal.

## Appendix B

### Simulation on the event width

In theory, any pattern defined in the sampling frequency of the data can be used for HMP analysis (see Section 2.1.1). In the case of EEG or MEG however, sine or half-sine waves are the expected patterns. As sine-waves can be decomposed into half-sines, these appear to be the minimal unit for an event in E/MEG. The expected duration of those half-sine templates for cognitive events is however unknown. In E/MEG, plausible durations include theta and alpha-band frequencies (Makeig, 2002; Yamanaka & Yamamoto, 2010). Anderson et al. (2016) used a 10Hz, hence 50ms duration half-sine. In this appendix, we illustrate the impact of target pattern duration on the estimations by HMP by simulating (as described in Section 3.1) three 25 or 100 ms events and trying to recover the ground truth with target pattern durations of 25, 50, 75, and 100 ms. The chosen source amplitude was 10^−6^*A* (SNR close to 10) to illustrate the general principle.

Figure B1 shows the resulting HMP solutions for 25 and 100ms generated pattern width. Overall it appears that the capture of the generated 25 ms events is successful for the 25ms and the 50ms target pattern (Figure B1a), but when searching for a 75ms or 100ms pattern one of the HMP event is missed and the timing of the correctly identified event is off when assuming too long target patterns (i.e. 100ms when simulating 25ms long patterns). When simulating a 100ms wide event to the contrary Figure B1b) we see that no matter the target pattern width all events are recovered, but when looking for much shorter events than the simulated durations (25ms target pattern in Figure B1), HMP estimates several events for a single simulated one.

To sum-up, without knowing the underlying duration of the events, expecting short event widths in HMP yields the best recovery performance, however, at the cost of increasing false positives. If researchers have expectations regarding duration of the events, then the best performance in terms of unique events will be achieved by aiming around the underlying true duration, the correct recovery of the events will however always be high as long as the expected duration is higher than the true underlying one.

## Appendix C

### Simulation on the probability distributions

For this appendix we manipulated the generative probability distribution as well as the average interval between events in the case of three sequential events. The simulation of HMP events was performed as described in Section 3.1 Because of the convolutions in Equation 5 and 6, if the events have a strong signal to noise ratio, the weight of the distribution will not strongly influence the result as long as credible values are not excluded (e.g., non-centered positive defined distributions). However, the expected distribution matters when signal-to-noise ratio is low. Therefore, we choose to simulate these datasets with a source amplitude of 10^−7^*A* (close to a SNR of 1 as in the right panel of Figure 3).

We simulated event times from the Weibull and the Wald distributions as those are often associated with reaction times (De Boeck & Jeon, 2019; Wagenmakers & Brown, 2007), which are directly related to event times. Additionally we also simulated from a gamma distribution as this was the original distribution used in (Anderson et al., 2016). For each of these distributions we selected 3 different shape values (Figure C1a). The choice of the shape values was made to represent a variety of moments and probability density spreads. For each of the 500 datasets, we first draw four average event times, for the three events and the response event, from a uniform distribution between 1 and 500 ms. We then simulate 100 trial event relative peak times from each of the 9 distributions with the drawn average between event interval (three families x three shapes). For a given dataset, we then try to recover the generative events using 9 HMP models all initialized with the selected distributions. As before we measured TPR and PPV (see Section A to test the performance of HMP.

As first observation, Figure C1b shows the TPR (color scale) and PPV (dot size) when data were generated with a given distribution (bottom row) and recovered with another or the same distribution (left most column). From this plot is can be seen that HMP performs best when the expected distribution corresponds to the generative one as shown by the higher TPR and PPV along the diagonal. It is also clear that the Wald and Gamma distributions seem to be most effective at recovering events generated with different distributions, and the Weibull distribution with a shape of 4 the worst.

In real data we do not know what generative distribution is underlying neural-time series in RT tasks. We therefore collapse HMP performance across all generative distributions and only look at the performances as a function of the expected distribution for HMP. As before, we use two binomial regression for both TPR and PPV on the logit scale, with the average reaction time (the sum of all distances between events), the expected distribution and the interaction between RT and distribution. The predictor of the average RT was centered and distributions were treated as a categorical factor with gamma 2 as a reference.

**Figure B1.**
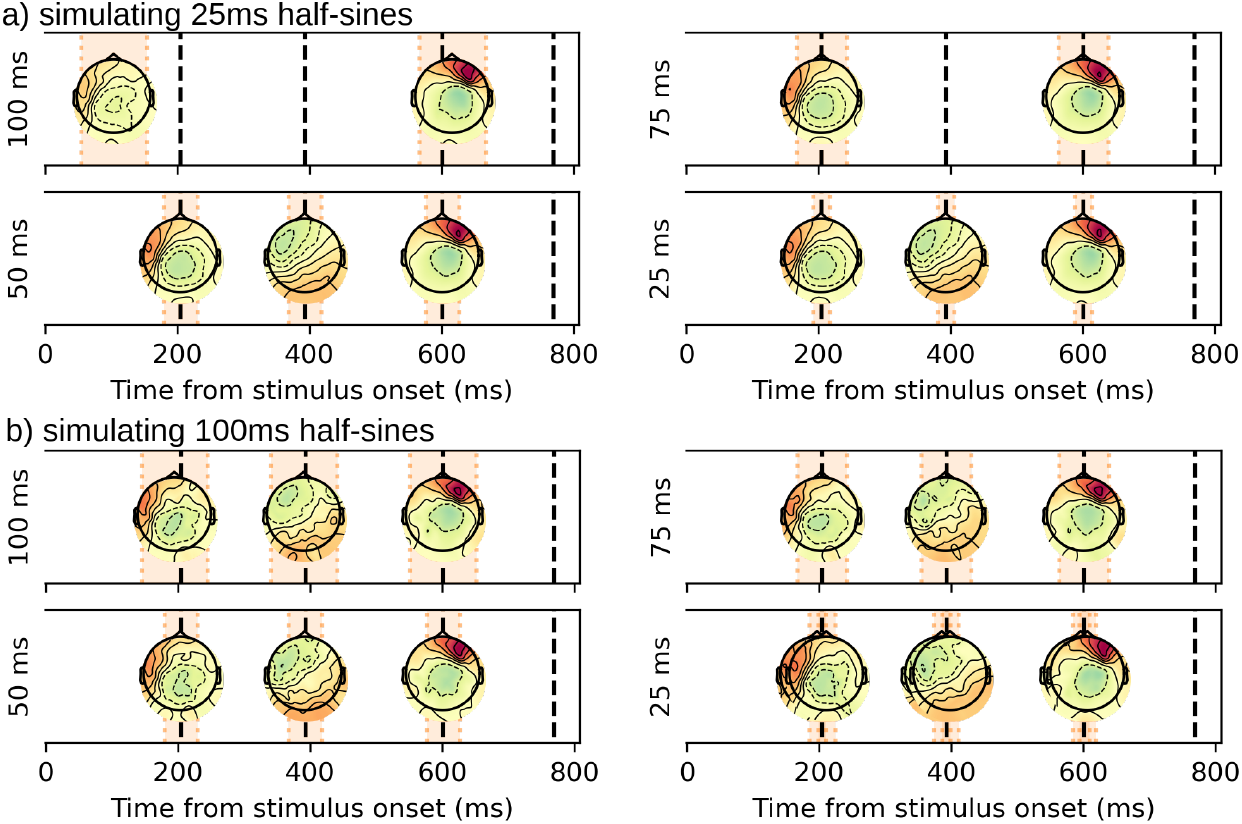
a) Example of solutions found when generating from a 25 ms event width for a target pattern of 100 ms, 75 ms, 50 ms or 25 ms. b) Example of solutions found when generating from a 100 ms event width for a target pattern of 100 ms, 75 ms, 50 ms or 25 ms.

The OSF repository contains all the detailed coefficients of the following linear models. For the TPR, the intercept (2.45, CrI = [2.39, 2.52]) shows that that the recovery of the true underlying events at the average RT for a gamma 2 is around 92%. This is comparable to the result we obtained in Simulation 1 with the same signal amplitude and distribution. When comparing the gamma 2 to other distributions, all other distributions, to the exception of the Wald-1 and the Weibull-4, have a slightly higher TPR (with the best performance for the Wald-0.25: 0.30, CrI = [0.20, 0.40]). For a worse performance on TPR some evidence is found for Wald with a shape of 1 (−0.31, CrI = [-0.40, -0.23]) and the Weibull with a shape of 4 (−0.55, CrI = [-0.63, -0.470]). The average RT has impact on the likelihood of recovering the true sequence for a gamma-2 (1.61, CrI = [1.34, 1.83]) predicting that adding one second the the average RT increases the TPR over 99% but shorter RTs are then associated with a much lower TPR. On the interaction between distribution and average RT, most interaction have strictly positive evidence while this means that the TPR is further improved for higher RTs (thus longer between event intervals) this also means that other distributions than the gamma 2 are performing worse on shorter event intervals (to the exception of the Wald 1 which is associated with a higher TPR for shorter event interval: -0.62, CrI = [-0.90, -0.31]).

The average PPV for a gamma-2 is around 83% (1.57, CrI = [1.52, 1.61]). All tested distributions show strictly negative evidence toward a lower performance than the gamma-2. The relation to the average RT is negative for the gamma-2 (−0.72, CrI = [-0.88, -0.57]). The evidence for a change in the relationship between RT and PPV for the other distributions is not found to the exception of the Wald-0.25 and the gamma-4 for which a weak evidence is found for an even lower PPV with increasing average RT.

In summary, compared to the gamma with a shape of 2 chosen by Anderson et al. (2016) other distributions, with some exceptions, perform very closely (i.e. the distance between lines in Figure C1). When some distributions perform statistically better, they also perform worse on the PPV. Based on these results, the original gamma 2 does seem like a good option, as it strikes a balance between identifying events while not overestimating the number of events. In general, the likelihood of recovering the sequence of events is dependent on the average RT (a proxy for the between-event interval) longer RTs predict better recovery performance but is also associated with increased false-alarms. Both of these effects are marginally modified for some distributions but again the overall performance is not drastically different for the tested distributions. Overall the choice of a distribution mostly boils down to where most of the probability mass should lie: if events are consistently very close in time to each other, a distribution with most probability mass towards lower value (e.g. a Wald-1) is preferred, while if they are consistently further appart, a more symmetric distribution would be preferred (e.g. Weibull-3).

**Figure C1.**
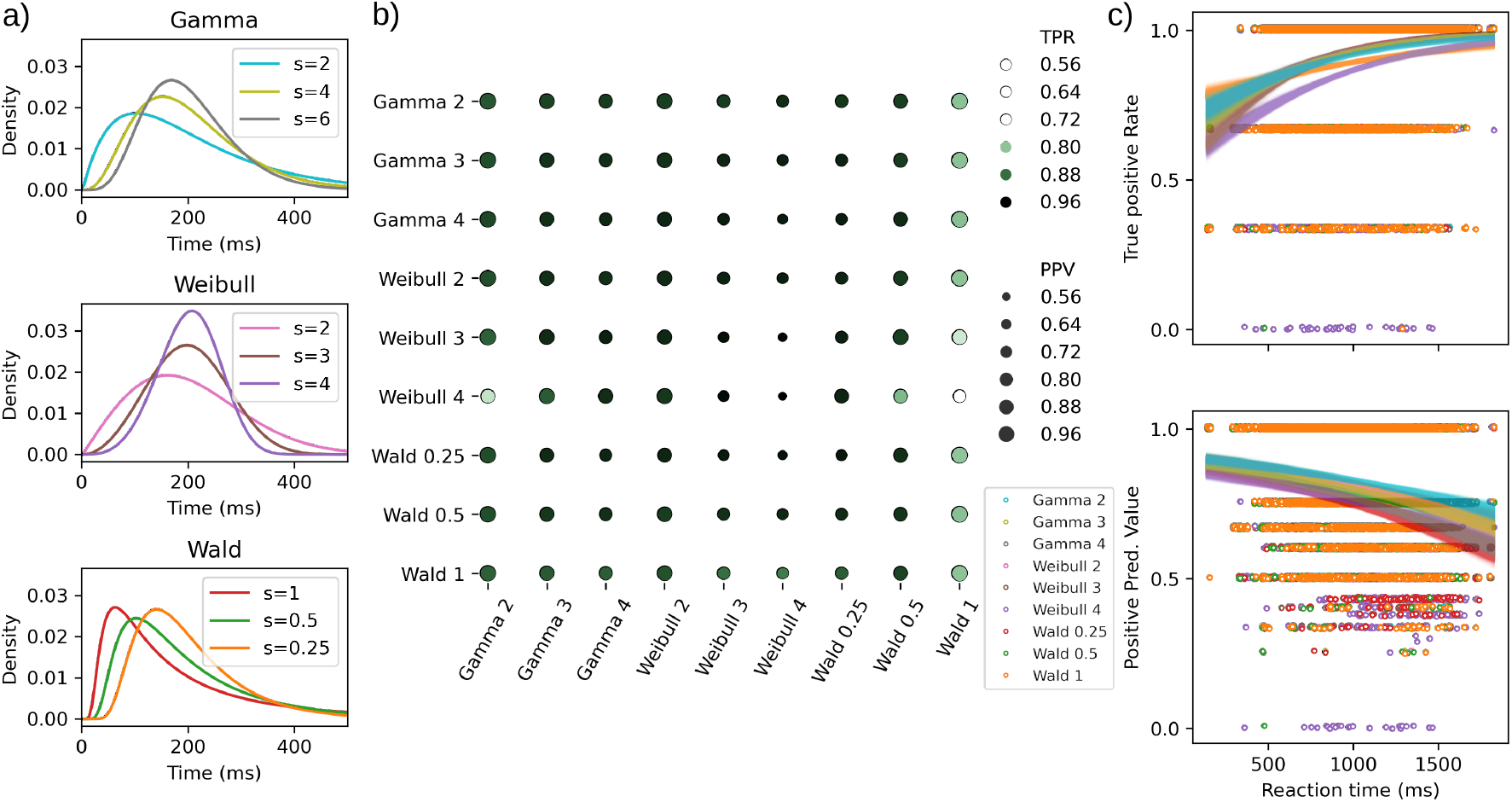
a) Representation of the different distribution used in the simulations, all have a mean of 200 ms for illustration purposes. b) Recovery matrix, the columns show the generative distribution while the row show the distribution used in HMP. TPR is represented by a color scale while PPV by the size of the dots. c) TPR (top) and PPV (bottom) for the 500 datasets with the different average RTs on the x-axis (ms) and colored by distribution used in HMP. Lines show the predicted values for 1000 draws from the posterior of each binomial regression.

## Footnotes

1 Note that this is not a required bandpass filter parameter to fit an HMP model, nor is it optimal to filter after epoching. This bandpass filtering is only used in the context of the replication

2 The solution resulting from HMP is not sensitive to the sampling rate in this case; downsampling to 100 Hz gives highly similar estimations.

3 This is not the case for model with more than one event as the event added after the first needs to be higher in log-likelihood than the one event solution.

## References

Anders, R., Alario, F., Van Maanen, L., et al. (2016). The shifted wald distribution for response time data analysis. Psychological methods, 21(3), 309.

Anderson, J. R., Borst, J. P., Fincham, J. M., Ghuman, A. S., Tenison, C., & Zhang, Q. (2018). The common time course of memory processes revealed. Psychological science, 29(9), 1463–1474.

Anderson, J. R., Zhang, Q., Borst, J. P., & Walsh, M. M. (2016). The discovery of processing stages: Extension of sternberg’s method. Psychological review, 123(5), 481.

Archambeau, K., Couto, J., & Van Maanen, L. (2023). Non-parametric mixture modeling of cognitive psychological data: A new method to disentangle hidden strategies. Behavior Research Methods, 55(5), 2232–2248.

Berberyan, H. S., van Maanen, L., van Rijn, H., & Borst, J. (2021). EEG-based Identification of Evidence Accumulation Stages in Decision-Making. Journal of Cognitive Neuroscience, 33(3), 510–527. 10.1162/jocn_a_01663

Berberyan, H. S., van Rijn, H., & Borst, J. P. (2021). Discovering the brain stages of lexical decision: Behavioral effects originate from a single neural decision process. Brain and Cognition, 153, 105786.

Boehm, U., Van Maanen, L., Forstmann, B., & van Rijn, H. (2014). Trial-by-trial fluctuations in cnv amplitude reflect anticipatory adjustment of response caution. NeuroImage, 96, 95–105.

Borst, J. P., & Anderson, J. R. (2024). Discovering cognitive stages in m/eeg data to inform cognitive models. In B. Forstmann & B. Turner (Eds.), An introduction to model-based cognitive neuroscience. Springer.

Botwinick, J., & Thompson, L. W. (1966). Premotor and motor components of reaction time. Journal of Experimental Psychology, 71(1), 9–15. 10.1037/h0022634

Brown, S., & Heathcote, A. (2005). A ballistic model of choice response time. Psychological Review, 112(1), 117–128. 10.1037/0033-295X.112.1.117

Brunet, D., Murray, M. M., & Michel, C. M. (2011). Spatiotemporal analysis of multichannel eeg: Cartool. Computational intelligence and neuroscience, 2011, 1–15.

Buc Calderon, C., Dewulf, M., Gevers, W., & Verguts, T. (2017). Continuous track paths reveal additive evidence integration in multistep decision making. Proceedings of the National Academy of Sciences, 114(40), 10618–10623. 10.1073/pnas.1710913114

Burle, B., Roger, C., Vidal, F., & Hasbroucq, T. (2008). Spatio-temporal dynamics of information processing in the brain: Recent advances, current limitations and future challenges. International Journal of Bioelectromagnetism, 10, 17–21.

Burle, B., Possamaï, C.-A., Vidal, F., Bonnet, M., & Hasbroucq, T. (2002). Executive control in the Simon effect: an electromyographic and distributional analysis. Psychological Research, 66(4), 324–36. 10.1007/s00426-002-0105-6

Burle, B., Vidal, F., Tandonnet, C., & Hasbroucq, T. (2004). Physiological evidence for response inhibition in choice reaction time tasks. Brain and cognition, 56(2), 153–164.

Cai, Y., Strauch, C., Van der Stigchel, S., & Naber, M. (2023). Open-dpsm: An open-source toolkit for modeling pupil size changes to dynamic visual inputs. Behavior Research Methods, 1–17.

Callaway, E., Halliday, R., Naylor, H., & Thouvenin, D. (1984). The latency of the average is not the average of the latencies. Psychophysiology, 21, 571.

Capretto, T., Piho, C., Kumar, R., Westfall, J., Yarkoni, T., & Martin, O. A. (2022). Bambi: A simple interface for fitting bayesian linear models in python. Journal of Statistical Software, 103(15), 1–29. 10.18637/jss.v103.i15

Christie, L. S., & Luce, R. D. (1956). Decision structure and time relations in simple choice behavior. The bulletin of mathematical biophysics, 18(2), 89–112.

Cohen, M. X. (2014). Analyzing neural time series data: Theory and practice. MIT press.

De Boeck, P., & Jeon, M. (2019). An overview of models for response times and processes in cognitive tests. Frontiers in psychology, 10, 102.

Donders, F. C. (1868). Die schnelligkeit psychischer processe: Erster artikel. Archiv für Anatomie, Physiologie und wissenschaftliche Medicin, 657–681.

Ehinger, B. V., & Dimigen, O. (2019). Unfold: An integrated toolbox for overlap correction, non-linear modeling, and regression-based eeg analysis. PeerJ, 7, e7838.

Eimer, M. (1996). The n2pc component as an indicator of attentional selectivity. Electroencephalography and clinical neurophysiology, 99(3), 225–234.

Forstmann, B. U., Wagenmakers, E.-J., Eichele, T., Brown, S., & Serences, J. T. (2011). Reciprocal relations between cognitive neuroscience and formal cognitive models: Opposites attract? Trends in cognitive sciences, 15(6), 272–279.

Froemer, R., Nassar, M. R., Ehinger, B. V., & Shenhav, A. (2022). Common neural choice signals emerge artifactually amidst multiple distinct value signals. BioRxiv, 2022–08.

Gramfort, A., Luessi, M., Larson, E., Engemann, D. A., Strohmeier, D., Brodbeck, C., Parkkonen, L., & Hämäläinen, M. S. (2014). Mne software for processing meg and eeg data. neuroimage, 86, 446–460.

Green, D. M. (1971). Fourier analysis of reaction time data. Behavior Research Methods & Instrumentation, 3(3), 121–125.

Groeneweg, E., Archambeau, K., & Van Maanen, L. (2021). Classification of cognitive strategies using EEG time series analyses. Proceedings of the 19th International Conference on Cognitive Modeling.

Jas, M., Engemann, D. A., Bekhti, Y., Raimondo, F., & Gramfort, A. (2017). Autoreject: Automated artifact rejection for meg and eeg data. NeuroImage, 159, 417–429.

Kappenman, E. S., Farrens, J. L., Zhang, W., Stewart, A. X., & Luck, S. J. (2021). Erp core: An open resource for human event-related potential research. NeuroImage, 225, 117465.

Kelly, S. P., & O’Connell, R. G. (2013). Internal and external influences on the rate of sensory evidence accumulation in the human brain. Journal of Neuroscience, 33(50), 19434–19441.

Kenemans, J., Baas, J., Mangun, G., Lijffijt, M., & Verbaten, M. (2000). On the processing of spatial frequencies as revealed by evoked-potential source modeling. Clinical neurophysiology, 111(6), 1113–1123.

Kirchherr, S., Mildiner Moraga, S., Coudé, G., Bimbi, M., Ferrari, P. F., Aarts, E., & Bonaiuto, J. J. (2023). Bayesian multilevel hidden markov models identify stable state dynamics in longitudinal recordings from macaque primary motor cortex. European Journal of Neuroscience, 58(3), 2787–2806.

Koenig, T., Prichep, L., Lehmann, D., Sosa, P. V., Braeker, E., Kleinlogel, H., Isenhart, R., & John, E. R. (2002). Millisecond by millisecond, year by year: Normative eeg microstates and developmental stages. Neuroimage, 16(1), 41–48.

Kruschke, J. K. (2010). Bayesian data analysis. Wiley Inter-disciplinary Reviews: Cognitive Science, 1(5), 658–676.

Kumar, R., Carroll, C., Hartikainen, A., & Martin, O. (2019). Arviz a unified library for exploratory analysis of bayesian models in python. Journal of Open Source Software, 4(33), 1143. 10.21105/joss.01143

Kutas, M., McCarthy, G., & Donchin, E. (1977). Augmenting mental chronometry: The p300 as a measure of stimulus evaluation time. Science, 197(4305), 792–795.

Lee, M. D., & Wagenmakers, E.-J. (2014). Bayesian cognitive modeling: A practical course. Cambridge university press.

Lehmann, D. (1971). Multichannel topography of human alpha eeg fields. Electroencephalography and clinical neurophysiology, 31(5), 439–449.

Luce, R. (1986). Response Times: Their Role in Inferring Elementary Mental Organization. Oxford University Press New York. 10.1093/acprof:oso/9780195070019.001.0001

Luck, S. J. (2005). Ten simple rules for designing and interpreting erp experiments. Event-related potentials: A methods handbook, 4.

Luck, S. J., & Hillyard, S. A. (1994). Electrophysiological correlates of feature analysis during visual search. Psychophysiology, 31(3), 291–308.

Makeig, S. (2002). Response: Event-related brain dynamics– unifying brain electrophysiology. Trends in neurosciences, 25(8), 390.

Masaracchia, L., Fredes, F., Woolrich, M. W., & Vidaurre, D. (2023). Dissecting unsupervised learning through hidden markov modelling in electrophysiological data. bioRxiv, 2023–01.

McClelland, J. L. (1979). On the time relations of mental processes: An examination of systems of processes in cascade. Psychological Review, 86, 287–330.

Meyer, D. E., Osman, A. M., Irwin, D. E., & Yantis, S. (1988). Modern mental chronometry. Biological Psychology, 26(1-3), 3–67.

Michel, C. M., & Koenig, T. (2018). Eeg microstates as a tool for studying the temporal dynamics of whole-brain neuronal networks: A review. Neuroimage, 180, 577–593.

Miller. (1993). A queue-series model for reaction time, with discrete-stage and continuous-flow models as special cases. Psychological Review, 100(4), 702.

Miller, J., Patterson, T., & Ulrich, R. (1998). Jackknife-based method for measuring lrp onset latency differences. Psychophysiology, 35(1), 99–115.

Miller, J., Ulrich, R., & Rinkenauer, G. (1999). Effects of stimulus intensity on the lateralized readiness potential. Journal of Experimental Psychology: Human Perception and Performance, 25(5), 1454.

Mouraux, A., & Iannetti, G. D. (2008). Across-trial averaging of event-related eeg responses and beyond. Magnetic resonance imaging, 26(7), 1041–1054.

Noorani, I., & Carpenter, R. (2016). The later model of reaction time and decision. Neuroscience & Biobehavioral Reviews, 64, 229–251.

Nunez, M. D., Gosai, A., Vandekerckhove, J., & Srinivasan, R. (2019). The latency of a visual evoked potential tracks the onset of decision making. Neuroimage, 197, 93–108.

O’connell, R. G., Dockree, P. M., & Kelly, S. P. (2012). A supramodal accumulation-to-bound signal that determines perceptual decisions in humans. Nature neuroscience, 15(12), 1729–1735.

Polich, J. (2007). Updating p300: An integrative theory of p3a and p3b. Clinical neurophysiology, 118(10), 2128–2148.

Portoles, O., Blesa, M., van Vugt, M., Cao, M., & Borst, J. P. (2022). Thalamic bursts modulate cortical synchrony locally to switch between states of global functional connectivity in a cognitive task. PLoS computational biology, 18(3), e1009407.

Posner, M. I. (1978). Chronometric Explorations of Mind. Lawrence Erlbaum.

Purcell, B. A., Heitz, R. P., Cohen, J. Y., Schall, J. D., Logan, G. D., & Palmeri, T. J. (2010). Neurally constrained modeling of perceptual decision making. Psychological review, 117(4), 1113.

Quinn, A. J., Vidaurre, D., Abeysuriya, R., Becker, R., Nobre, A. C., & Woolrich, M. W. (2018). Task-evoked dynamic network analysis through hidden markov modeling. Frontiers in neuroscience, 12, 603.

Rabiner, L. R. (1989). A tutorial on hidden markov models and selected applications in speech recognition. Proceedings of the IEEE, 77(2), 257–286.

Ratcliff, R. (1978). A theory of memory retrieval. Psychological Review, 85(2), 59–108. 10.1037/0033-295X.85.2.59

Selen, L. P. J., Shadlen, M. N., & Wolpert, D. M. (2012). Deliberation in the Motor System: Reflex Gains Track Evolving Evidence Leading to a Decision. Journal of Neuroscience, 32(7), 2276–2286. 10.1523/JNEUROSCI.5273-11.2012

Servant, M., Logan, G. D., Gajdos, T., & Evans, N. J. (2021). An integrated theory of deciding and acting. Journal of Experimental Psychology: General, 150(12), 2435.

Smith, P. L. (1990). Obtaining meaningful results from fourier deconvolution of reaction time data. Psychological Bulletin, 108(3), 533.

Smith, P. L. (1995). Psychophysically principled models of visual simple reaction time. Psychological Review, 102(3), 567–593. 10.1037/0033-295X.102.3.567

Smulders, F. T., Kenemans, J., & Kok, A. (1994). A comparison of different methods for estimating singletrial p300 latencies. Electroencephalography and Clinical Neurophysiology/Evoked Potentials Section, 92(2), 107–114.

Stone, M. (1960). Models for choice-reaction time. Psychometrika, 25(3), 251–260. 10.1007/BF02289729

Tenison, C., Fincham, J. M., & Anderson, J. R. (2016). Phases of learning: How skill acquisition impacts cognitive processing. Cognitive psychology, 87, 1–28.

Thura, D., Cabana, J.-F., Feghaly, A., & Cisek, P. (2022). Integrated neural dynamics of sensorimotor decisions and actions. PLoS biology, 20(12), e3001861.

Townsend, J. T., & Nozawa, G. (1995). Spatio-temporal properties of elementary perception: An investigation of parallel, serial, and coactive theories. Journal of Mathematical Psychology, 39(4), 321–359.

Turner, B. M., Forstmann, B. U., Love, B. C., Palmeri, T. J., & Van Maanen, L. (2017). Approaches to analysis in model-based cognitive neuroscience. Journal of Mathematical Psychology, 76, 65–79.

Van de Ville, D., Britz, J., & Michel, C. M. (2010). Eeg microstate sequences in healthy humans at rest reveal scale-free dynamics. Proceedings of the National Academy of Sciences, 107(42), 18179–18184.

van der Velde, M., Sense, F., Borst, J. P., Van Maanen, L., & Van Rijn, H. (2022). Capturing dynamic performance in a cognitive model: Estimating act-r memory parameters with the linear ballistic accumulator. Topics in Cognitive Science, 14(4), 889–903.

Van Maanen, L., de Jong, R., & van Rijn, H. (2014). How to assess the existence of competing strategies in cognitive tasks: A primer on the fixed-point property. PloS one, 9(8), e106113.

Van Maanen, L., Portoles, O., & Borst, J. P. (2021). The discovery and interpretation of evidence accumulation stages. Computational brain & behavior, 4(4), 395–415.

Van Maanen, L., van Rijn, H., & Borst, J. P. (2009). Stroop and picture—word interference are two sides of the same coin. Psychonomic bulletin & review, 16, 987–999.

Visser, I. (2011). Seven things to remember about hidden markov models: A tutorial on markovian models for time series. Journal of Mathematical Psychology, 55(6), 403–415.

von Helmholtz, H. (1850). Mittheilung für die physikalische gesellschaft in berlin betreffend versuche über die fortpflanzungsgeschwindigkeit der reizung in den sensiblen nerven des menschen. Archive of the BerlinBrandenburgische Akademie der Wissenschaften, 1–4.

Wagenmakers, E.-J., & Brown, S. (2007). On the linear relation between the mean and the standard deviation of a response time distribution. Psychological review, 114(3),830.

Weindel, G., Anders, R., Alario, F., Burle, B., et al. (2021). Assessing model-based inferences in decision making with single-trial response time decomposition. Journal of Experimental Psychology: General, 150(8), 1528.

Weindel, G., Gajdos, T., Burle, B., & Alario, F.-X. (2022). The decisive role of non-decision time for interpreting decision making models. PsyArXiv.

Woody, C. D. (1967). Characterization of an adaptive filter for the analysis of variable latency neuroelectric signals. Medical and biological engineering, 5, 539–554.

Wu, C. J. (1983). On the convergence properties of the em algorithm. The Annals of statistics, 95–103.

Yamanaka, K., & Yamamoto, Y. (2010). Single-trial eeg power and phase dynamics associated with voluntary response inhibition. Journal of Cognitive Neuroscience, 22(4), 714–727.

Yu, S.-Z. (2010). Hidden semi-markov models. Artificial intelligence, 174(2), 215–243.

Zhang, Q., Van Vugt, M., Borst, J. P., & Anderson, J. R. (2018). Mapping working memory retrieval in space and in time: A combined electroencephalography and electrocorticography approach. NeuroImage, 174, 472–484. 10.1016/j.neuroimage.2018.03.039

